# Genetically encoded sensors for measuring histamine release both *in vitro* and *in vivo*

**DOI:** 10.1101/2022.08.19.504485

**Authors:** Hui Dong, Mengyao Li, Yuqi Yan, Tongrui Qian, Yunzhi Lin, Xiaoyuan Ma, Henry F. Vischer, Can Liu, Guochuan Li, Huan Wang, Rob Leurs, Yulong Li

## Abstract

Histamine (HA) is a key biogenic monoamine involved in a wide range of physiological and pathological processes in both the central nervous system and the periphery. Because the ability to directly measure extracellular HA in real-time will provide important insights into the functional role of HA in complex circuits under a variety of conditions, we developed a series of genetically encoded <u>G</u> protein-coupled <u>r</u>eceptor <u>a</u>ctivation-<u>b</u>ased (GRAB) HA (GRAB_HA_) sensors. These sensors produce a robust increase in fluorescence upon HA application, with good photostability, sub-second kinetics, nanomolar affinity, and high specificity. Using these GRAB_HA_ sensors, we measured electrical stimulation-evoked HA release in acute brain slices with high spatiotemporal resolution. Moreover, we recorded HA release in the preoptic area of the hypothalamus and in the medial prefrontal cortex during the sleep-wake cycle in freely moving mice, finding distinct patterns of HA release in these specific brain regions. Together, these *in vitro* and *in vivo* results show that our GRAB_HA_ sensors have high sensitivity and specificity for measuring extracellular HA, thus providing a robust new set of tools for examining the role of HA signaling in both health and disease.

## Introduction

The biogenic amine histamine (HA) is an important signaling molecule in the immune, digestive, and nervous systems. For example, HA is secreted by basophils and mast cells as part of a localized immune response, playing a role in allergy and itch^1^. HA is also released by enterochromaffin-like cells in the stomach, triggering the release of stomach acids^2^. Since the discovery that antihistamines also have sedative properties, the role of HA in the central nervous system has attracted considerable attention^3^. In the vertebrate brain, HA is synthesized primarily in the tuberomammillary nucleus (TMN) in the posterior hypothalamus^4^, and histaminergic TMN neurons project throughout the brain, regulating various functions such as the sleep-wake cycle, feeding, attention, and learning^5^.

Despite its clearly important role in a wide range of physiological and pathological processes, the spatiotemporal dynamics of HA release during various behaviors remain poorly understood, due in large part to the limitations associated with current detection methods. For example, microdialysis combined with analytical techniques has been widely used to measure the dynamics of HA release in the living brain^6^. However, microdialysis often has low temporal resolution due to the relatively long time needed to collect the samples. In addition, complementary approaches using electrochemical methods such as fast-scan cyclic voltammetry have been developed for detecting HA in real time^7^. However, both microdialysis and electrochemical methods have limited spatial precision and lack cell-type specificity. Coupling diamine oxidase with an optical oxygen nanosensor can provide continuous tracking of HA concentration^8^, but the nanosensor has low sensitivity (with a lower limit of detection of approximately 1.1 mM) and specificity, and is less effective at detecting HA in deep brain regions. The Tango-Trace strategy can be used to detect HA release *in vivo* with a high signal-to-background ratio by coupling the β-arrestin signaling pathway to the expression of a reporter gene^9–11^. However, this approach requires several hours to express the fluorescent reporter protein and cannot be used to monitor the rapid dynamics of HA-mediated transmission. Finally, conformational H_1_R and H_3_R sensors based on both fluorescence resonance energy transfer (FRET) and bioluminescence resonance energy transfer (BRET) have been developed to measure their interaction with histamine (and mechanical force or other H_3_R ligands, respectively)^12–14^, but generally have a low signal-to-noise ratio and a low dynamic range, thus strongly limiting their ability to monitor HA release *in vivo*.

Recently, building on the successful G protein-coupled receptor activation–based (GRAB) strategy, our group and others independently developed a series of genetically encoded sensors for detecting a variety of neurotransmitters and neuromodulators with high sensitivity, selectivity, and spatiotemporal resolution in *in vivo* preparations^15–19^. Using this strategy, we developed a pair of genetically encoded fluorescent sensors called GRAB_HA1h_ and GRAB_HA1m_ (abbreviated here as HA1h and HA1m, respectively) based on the human H_4_R and water bear (tardigrade) H_1_R receptors, respectively, in order to measure extracellular HA with high sensitivity and high spatiotemporal resolution both *in vitro* and *in vivo*. These sensors have high specificity for HA, rapid kinetics (on the order of sub-second), and an increase in fluorescence of approximately 300-500% in response to HA when applied *in vitro*. Using these novel HA sensors, we monitored the release of endogenous HA during the sleep-wake cycle in freely moving mice and found distinct patterns of HA release in two specific brain regions. Thus, these genetically encoded sensors can be used to gain important new insights into the dynamic properties of HA signaling under both physiological and pathological conditions.

## Results

### Development and characterization of GRAB sensors for detecting histamine

To monitor the change in extracellular HA levels with high spatial and temporal resolution, we designed a genetically encoded GRAB sensor based on HA receptors (Fig. 1a). First, we systematically screened a series of G protein-coupled HA receptors, including human H_1_R, H_2_R, H_3_R, and H_4_R (Supplementary Fig. 1a,b). The third intracellular loop (ICL3) in the HA receptors was replaced with a circularly permutated EGFP (cpEGFP) and ICL3 derived from the previously developed and characterized GRAB_NE_ norepinephrine sensor^20^. We selected the hH_4_R-based chimera, which has good plasma membrane trafficking (Supplementary Fig. 1b), for further optimization. We then optimized the length and amino acid composition of the linker region near the insertion site, as well as critical residues in cpEGFP based on our previous experience in developing GRAB sensors (Fig. 1b). Screening more than 2,000 sensor variants resulted in the sensor with the highest fluorescence response, which we called HA1h (Fig. 1b, Supplementary Fig. 2a). When expressed in cultured HEK293T cells, HA1h trafficked to the plasma membrane and produced a peak change in fluorescence (ΔF/F_0_) of ~370% in response to extracellular application of 10 μM HA (Fig. 1f-i). We also generated an HA-insensitive form of this sensor, which we call HA1mut, by introducing an E211A substitution in hH_4_R (Fig. 1b, Supplementary Fig. 2a). HA application induced a dose-dependent increase in fluorescence in HA1h-expressing HEK293T cells, with a half-maximal effective concentration (EC50) of 17 nM (Fig. 1j). Consistently, a similar binding affinity (K_d_) of 13 nM was obtained for the binding of [^3^H]HA to HA1h in a radioligand binding assay (Supplementary Fig. 5a). We also sought to develop a more sensitive HA sensor with a higher response and a larger dynamic range by screening HA receptors obtained from 11 species in which we grafted the ICL3-cpEGFP module from the HA1h sensor (Fig. 1c,d). Based on the results of this screen, we selected the sensor based on the water bear (tardigrade) H_1_R for further optimization, yielding a high-response sensor we call HA1m (Fig. 1e, Supplementary Fig. 2b), which produces an ~590% increase in fluorescence upon application of 100 μM HA (Fig. 1f-i) and an EC50 of 380 nM when expressed in HEK293T cells (Fig. 1j).

**Fig. 1:**
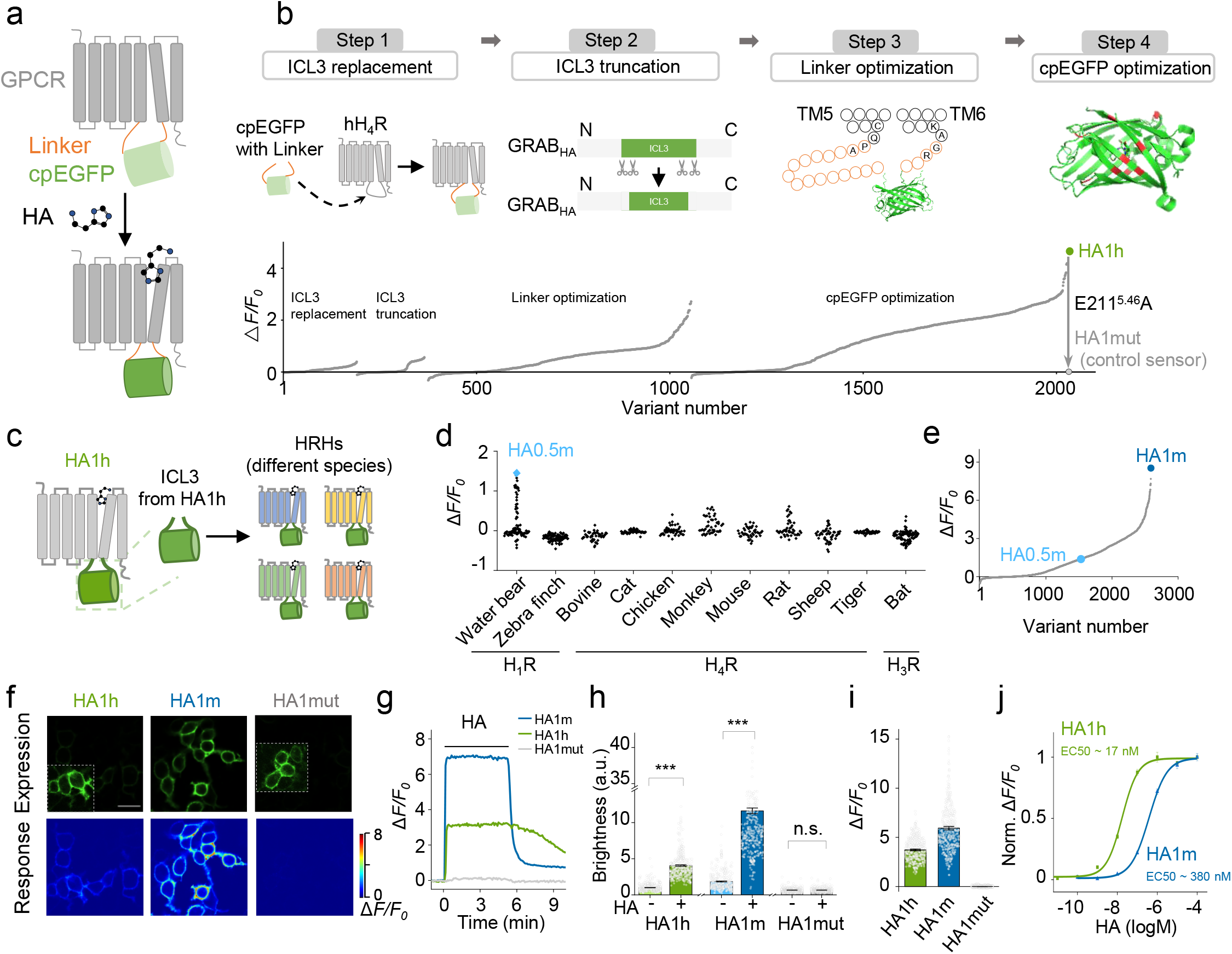
Development of genetically encoded fluorescent GRAB_HA_ sensors. (a) Schematic depiction of the principle behind the GRAB_HA_ sensors in which cpEGFP is inserted within the receptor. Upon binding HA, the resulting conformational change causes an increase in fluorescence. (b) Steps used to identify the most responsive candidate HA sensors based on the human H_4_R and GRAB_NE_ ICL3, by screening the ICL3 insertion sites, the ICL3 linker length, and key residues in the linker and cpEGFP. Shown is the final HA1h sensor used in this study, as well as the corresponding HA1mut sensor used in this study as a negative control. (c) Schematic depiction of the principle used to generate and screen GRAB_HA_ sensors derived from other species’ HA receptors. (d) Further development of HA sensors by screening H1, H3, and H4 receptors obtained from the indicated species. (e) Optimization of candidate HA sensors based on the water bear (tardigrade) H_1_R. (f-j) Characterization of the expression and performance of the HA1h, HA1m, and HA1mut sensors expressed in HEK293T cells, showing membrane trafficking (f), representative time courses (g), relative brightness (h), peak response to HA (i), and apparent affinity (j). Scale bar, 50 μm. Data are shown as mean ± s.e.m. in h,i,j with the error bars or shaded regions indicating the s.e.m., *** *P* < 0.001; n.s., not significant.

We then characterized the kinetic properties and wavelength spectrum of the HA sensors. Using a local perfusion system and high-speed line-scanning, both HA1h and HA1m had a rapid increase in fluorescence in response to a saturating concentration of HA, with an on-rate (time constant) of 0.6 s and 0.3 s, respectively (Fig. 2a-c). The off-rate of HA1h and HA1m was measured by rapidly applying H_4_R and H_1_R antagonists, respectively, yielding an off-rate (time constant) of 2.3 s (HA1m) and 1.4 s (HA1m) (Fig. 2a-c). Both HA1h and HA1m had a similar spectrum as EGFP, with an excitation peak at ~505 nm and an emission peak at ~520 nm under one-photon illumination, and an excitation peak at ~920 nm under two-photon illumination (Supplementary Fig. 3a).

**Fig. 2:**
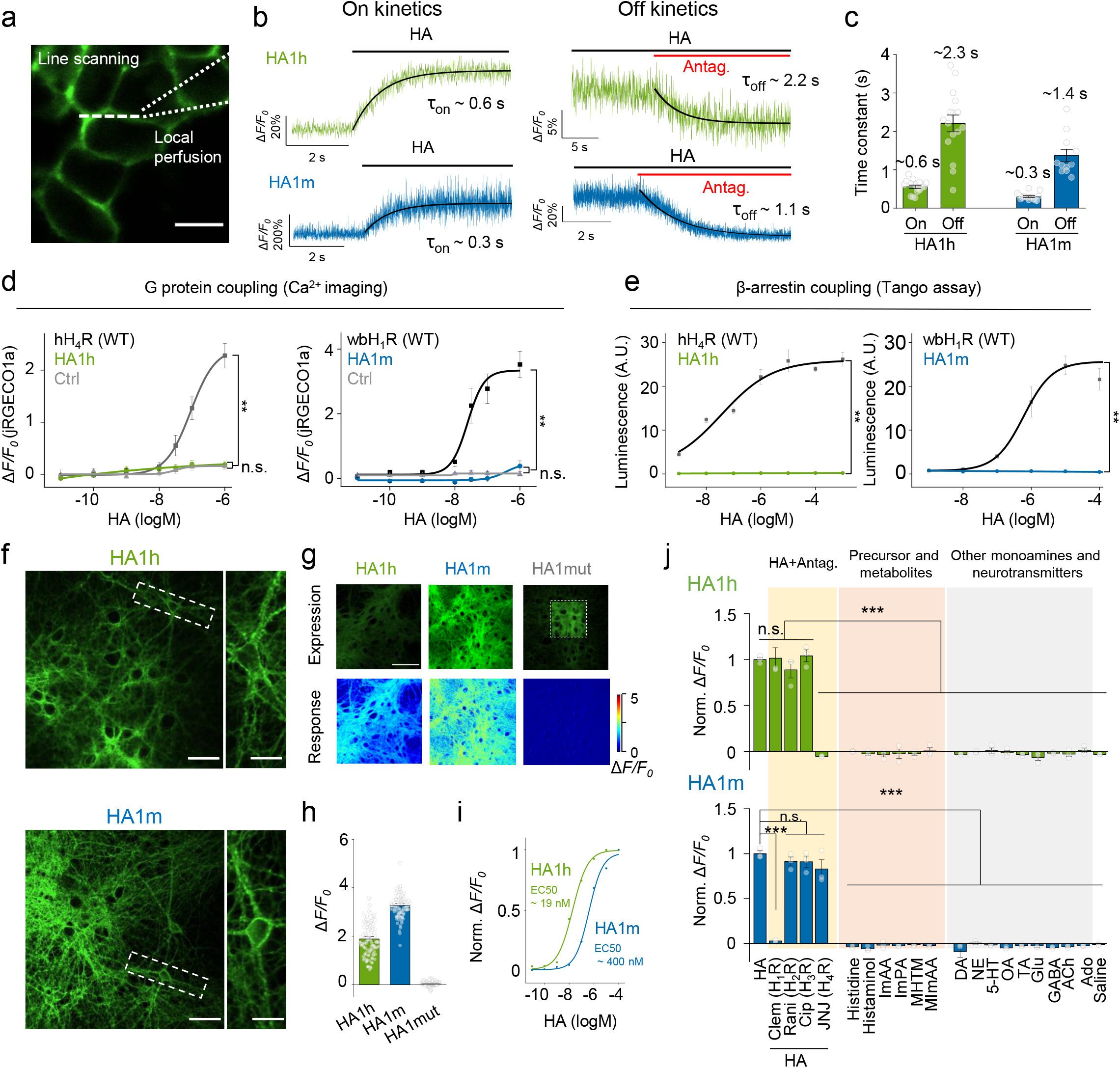
Characterization of GRAB_HA_ sensors in HEK293T cells and cultured rat cortical neurons. (a-c) Measurement of the kinetics of HA1h and HA1m expressed in HEK293T cells. HA was applied to cells expressing either HA1h or HA1m in order to measure τ_on_. The appropriate antagonist (100 μM JNJ-7777120 for HA1h, 100 μM clemastine for HA1m) was then applied in the continued presence of HA in order to measure τ_off_. Representative traces (b) and summary results (c) are shown; the white dashed line in (a) indicates the line-scanning region. *n* ≥ 10 cells from 4-5 cultures. Scale bar, 50 μm. (d and e) G protein (d) and β-arrestin (e) coupling were measured in cells expressing HA1h (or H_4_R as a positive control) or HA1m (or H_1_R as a positive control) using Ca^2+^ imaging (d) and a Tango assay (e). ** *P* < 0.01; n.s., not significant. (f-i) Characterization of the expression and performance of the HA1h, HA1m, and HA1mut sensors expressed in cultured rat cortical neurons, showing membrane trafficking (f), representative responses (g), peak response to HA (h), and apparent affinity (i). Scale bar, 100 μm left panel, 20 μm right panel in f; 100 μm in g. (j) Summary of the normalized fluorescence change measured in cultured rat cortical neurons expressing HA1h (top panel) or HA1m (bottom panel) in response to HA alone, HA applied together with the indicated HA receptor antagonist, HA precursor, metabolites, and the indicated neurotransmitters (all compounds were applied at 5 μM). Clem: clemastine; Rani: ranitidine; Cip: ciproxifan; JNJ: JNJ-7777120; ImAA: imidazoleacetic acid; ImPA: imidazolepyruvic acid; MHTM: 1-methylhistamine; MImAA: methylimizoleacetic acid; DA: dopamine; NE: norepinephrine; 5-HT: serotonin; OA: octopamine; TA: tyramine; Glu: glutamate; GABA: γ-aminobutyric acid; ACh: acetylcholine; Ado: adenosine. *n* = 3 wells for different groups. Paired two-tailed Student’s t-tests were performed. ** *P* < 0.01; n.s., not significant. Data are shown as mean ± s.e.m. in c,d,e,h,i,j with the error bars indicating the s.e.m.

Next, we tested whether our HA sensors can trigger downstream signaling pathways, including G protein–dependent and/or β-arrestin–dependent pathways. Because human H_4_R is a Gi-coupled G protein–coupled receptor (GPCR), we used a chimeric Gα_q-i_ protein, which switches the downstream response from Gi to Gq signaling^21,22^. Using intracellular Ca^2+^ imaging, we found that the hH_4_Rs and water bear H_1_Rs signaled robustly, whereas both the HA1h and HA1m sensors produced virtually no response (Fig. 2d). Similar results were obtained when we measured signaling via the β-arrestin pathway using the Tango assay (Fig. 2e). Furthermore, consistent with our Tango assay results, we found that exposing HA1h- and HA1m-expressing cultured neurons to a saturating concentration of HA for 3 h led to virtually no internalization of either sensor (Supplementary Fig. 4a,b). Taken together, these results suggest that our HA sensors can be used to measure extracellular HA without inadvertently activating downstream signaling pathways.

To measure the performance of our HA sensors in a more physiologically relevant context, we expressed HA1h, HA1m, or HA1mut in cultured rat cortical neurons using an adeno-associated virus (AAV). We found that HA1h and HA1m localize throughout the neuronal plasma membrane (Fig. 2f). Moreover, bath application of a saturating concentration of HA caused a robust increase in both HA1h and HA1m fluorescence (with peak ΔF/F_0_ values of approximately 180% and 320%, respectively), with no measurable effect on HA1mut (Fig. 2g,h). In addition, similar to our results obtained with HEK293T cells, we found that neurons expressing either HA1h or HA1m had a dose-dependent increase in fluorescence, with EC50 values of 19 nM and 400 nM, respectively (Fig. 2i). To confirm the specificity of our sensors for HA, we tested the effects of an HA precursor, several HA metabolites, and several monoamines and neurotransmitters, finding no measurable response to any of these compounds (Fig. 2j). Notably, the HA-induced response in HA1h and HA1m was fully blocked by the H_4_R antagonist JNJ-7777120 and the H_1_R antagonist clemastine, respectively, while all other HA receptor antagonists had no effect (Fig. 2j). Consistently, the HA-induced response in HA1h and HA1m was specifically blocked by the H_4_R antagonist and the H_1_R antagonist in cultured HEK293T cell systems (Supplementary Fig. 3b), respectively. Furthermore, radioligand binding assay was performed to test whether HA sensors specifically are bound by HA. We found HA and the H_4_R agonist *N*^α^-methylhistamine to specifically bind to HA1h sensor, while HA precursor and HA metabolites, and several monoamines and neurotransmitters had no measurable binding (Supplementary Fig. 5b). Together, these results indicate that both HA1h and HA1m retain the pharmacological profile and ligand specificity of their corresponding parent receptors.

### GRAB_HA_ sensors can be used to image HA release in acute mouse brain slices

Next, we examined whether our HA sensors can be used to detect the release of endogenous HA. We therefore injected an AAV expressing HA1m under the control of the human synapsin (hSyn) promoter into the mouse prefrontal cortex (PFC) and prepared acute brain slices 2-3 weeks later (Fig. 3a,b). We found that applying electrical stimuli at 20 Hz elicited a pulse number–dependent increase in HA1m fluorescence that was blocked by treating the slices with the H_1_R antagonist clemastine (Fig. 3c,d). We also measured the kinetics of the response to 10, 20, 50, and 100 pulses delivered at 20 Hz and obtained rise time constants (τ_on_) and decay time constants (τ_off_) of 1.4-5.2 s and 10-13 s, respectively (Fig. 3e). Interestingly, we also found that the HA1m signal increased at the site of stimulation and then propagated outward (Fig. 3f). We measured the change in fluorescence at various distances from stimulation site at various time points and obtained an apparent diffusion coefficient of approximately 1.9 x 10^3^ μm^2^/s (Fig. 3g-j). Thus, our HA1m sensor can detect the release of endogenous HA with high spatiotemporal resolution.

**Fig. 3:**
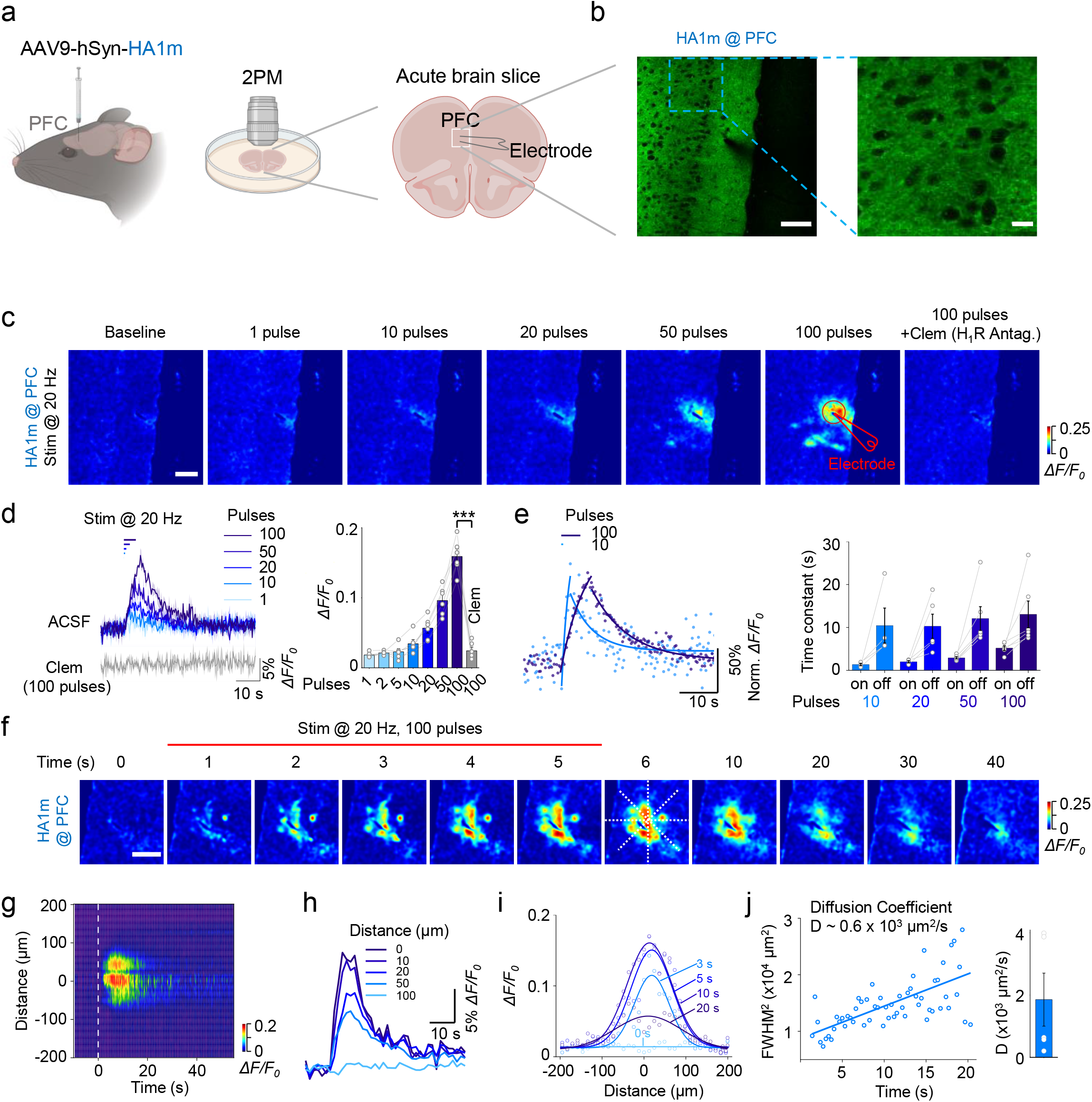
Using the GRAB_HA_ sensor to measure the release of endogenous HA in acute mouse brain slices. (a) Schematic illustration of the acute mouse brain slice experiments. An AAV expressing hSyn-HA1m was injected into the PFC region; acute brain slices (coronal slices containing the PFC) were then prepared and used for electrical stimulation experiments. (b) Exemplar 2-photon microscopy images showing HA1m expressed in the PFC. Scale bars: 100 μm (left) and 20 μm (right). (c) Representative pseudocolor images showing the corresponding fluorescence change in HA1m-expressing slices in response to 1, 10, 20, 50, and 100 pulses delivered at 20 Hz in ACSF, and in response to 100 pulses in the presence of the H_1_R antagonist clemastine (20 μM). Scale bar, 100 μm. (d) Exemplar traces (left) and summary of the peak fluorescence change (right) measured in HA1m-expressing slices stimulated as indicated. Where indicated, clemastine was applied at 20 μM. *** *P* < 0.001. (e) Representative trace with fitted curves (left) and summary data of the on-time and off time constants (right) measured for the change in HA1m fluorescence in response to the indicated number of pulses applied at 20 Hz. (f) Exemplar time-lapse pseudocolor images of HA1m fluorescence in response to 100 pulses at 20 Hz applied during the first 5 s. Scale bar, 100 μm (g) Spatial profile of the evoked change in fluorescence shown in panel (f); the vertical dashed line at time 0 indicates the start of the stimulation. (h) Temporal dynamics of the change in HA1m fluorescence as shown in (g) measured 10, 20, 50, and 100 μm from the release center. (i) Spatial dynamics of the change in HA1m fluorescence as shown in (g) measured 3, 5, 10, and 30 s after the onset of stimulation. Each curve was fitted with a Gaussian function. (j) Left, a representative diffusion coefficient was measured by plotting the square of the full width at half maximum (FWHM) against time based on the data show in (i). Right, summary of the diffusion coefficients measured from the data shown in (i). Data are shown as mean ± s.e.m. in d,e,j with the error bars indicating the s.e.m.

### GRAB_HA_ sensors can be used to measure HA release *in vivo*

A wealth of pharmacological and genetic data has shown that HA plays an essential role in regulating the sleep-wake cycle^23^. Thus, based on these findings we examined whether our HA sensors could be used to measure histaminergic activity during the physiological sleep-wake cycle, focusing on the transitions between sleep-wake states. In the mammalian brain, the preoptic area (POA) regulates sleep and receives extensive projections from histaminergic neurons^24,25^. We expressed the HA1m sensor in the POA and then performed simultaneous fiber photometry, electroencephalography (EEG), and electromyography (EMG) recordings in freely behaving mice (Fig. 4a). We found that the signal produced by the HA1m sensor in the POA was relatively high when the mouse was awake compared to the signal measured during both rapid eye movement (REM) and non-REM (NREM) sleep (Fig. 4b,c), supporting previously reported data obtained using microdialysis^26^. As a control, we also expressed the HA1m sensor in the POA of histidine decarboxylase knockout (HDC KO) mice and found no significant change in HA1m fluorescence at any stage during the sleep-wake cycle (Fig. 4d). Because our HA sensors have rapid kinetics, we then calculated the change in HA1m signal at the transitions between various sleep states and found that the signal increased during the REM→wake and NREM→wake transitions, but decreased during the wake→NREM and NREM→REM transitions (Fig. 4f).

**Fig. 4:**
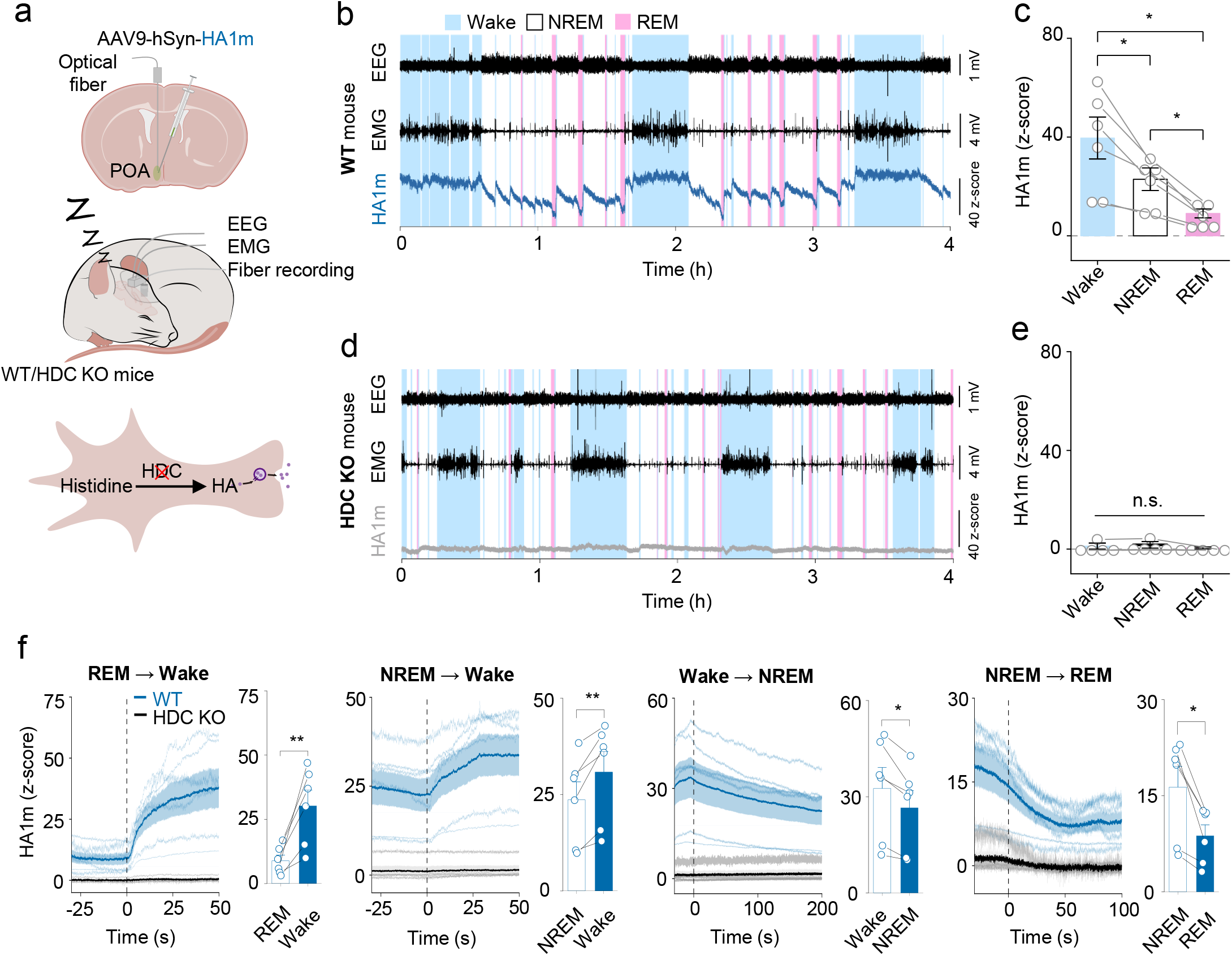
Using the GRABHA1m sensor to measure *in vivo* HA release in the POA during the sleep-wake cycle in freely moving mice. (a) Schematic diagram depicting the strategy for fiber photometry recording of HA1m fluorescence in the preoptic area (POA) of freely behaving wild-type (WT) and histidine decarboxylase knockout (HDC KO) mice during the sleep-wake cycle. Top panel, an AAV expressing hSyn-HA1m was injected into the PFC region; middle panel, schematic diagram depicting fiber photometry recording of HA1m signals, EEG, and EMG. Bottom panel, HA synthesis process. (b and d) Example traces of simultaneous EEG, EMG, and HA1m recordings in a WT (b) and HDC KO (d) mouse. In this and subsequent figures, the wake state is shaded in blue, REM sleep is shaded in pink, and NREM sleep is unshaded. (c and e) Summary of the average HA1m fluorescence measured in WT (c; n=6 mice) and HDC KO mice (e; *n* = 5 mice) in the indicated sleep-wake states. Each symbol represents the recording from one mouse. (c) One-way ANOVA: *F* = 16.3913, *P* = 0.0007; Tukey’s *post hoc* test: Wake vs NREM *P* = 0.0335, Wake vs REM *P* = 0.0207, NREM vs REM *P* = 0.0153; * *P* < 0.05. (e) One-way ANOVA: *F* = 1.2401, *P* = 0.3308; n.s., not significant. (f) Time course and average signals measured for the HA1m sensor during the indicated transitions between sleep-wake states. The vertical dashed lines at time 0 represent the transition time. paired t-test, REM→Wake *t* = 4.2803, *P* = 0.0079; NREM→Wake *t* = 4.0586, *P* = 0.0097; Wake→NREM *t* = 3.7418, *P* = 0.0134; NREM→REM *t* = 3.9835, *P* = 0.0105; * *P* < 0.05, ** *P* < 0.01. Data are shown as mean ± s.e.m. in c,e,f with the error bars or shaded regions indicating the s.e.m. in f, the shaded lines indicating the mean trace of individual mice.

We recorded HA release in the POA using our medium-affinity HA1m sensor, as the POA receives a relatively dense number of histaminergic projections. Next, we examined whether we could measure HA release in the prefrontal cortex (PFC), a brain region that receives a more moderate density of histaminergic projections and plays a critical role in regulating executive functions such as attention and decision, functions that are based on wakefulness. To account for the lower density of histaminergic projections, we expressed the high-affinity HA1h sensor in the PFC and performed simultaneous fiber photometry, EEG, and EMG recordings in freely behaving mice (Fig. 5a). Similar to our results obtained using the HA1m sensor expressed in the POA, we found that the HA1h signal in the PFC was higher when the animal was awake compared to during both REM and NREM sleep (Fig. 5b,c), again consistent with previously reported microdialysis data ^27^. As expected, consistent with our *in vitro* results, an i.p. injection of the H_4_R antagonist JNJ-7777120 virtually eliminated fluorescence of the HA1h sensor expressed in the PCF, even when the mouse was awake (Fig. 5d,e). Furthermore, no significant change in fluorescence was measured in the PFC of mice expressing the mutant HA1mut sensor at any time during the sleep-wake cycle (Fig. 5f,g). Finally, no significant attenuation in HA1h sensor signals over one day recording under low excitation light intensity and sample rate (Supplementary Fig. 7a-c).

**Fig. 5:**
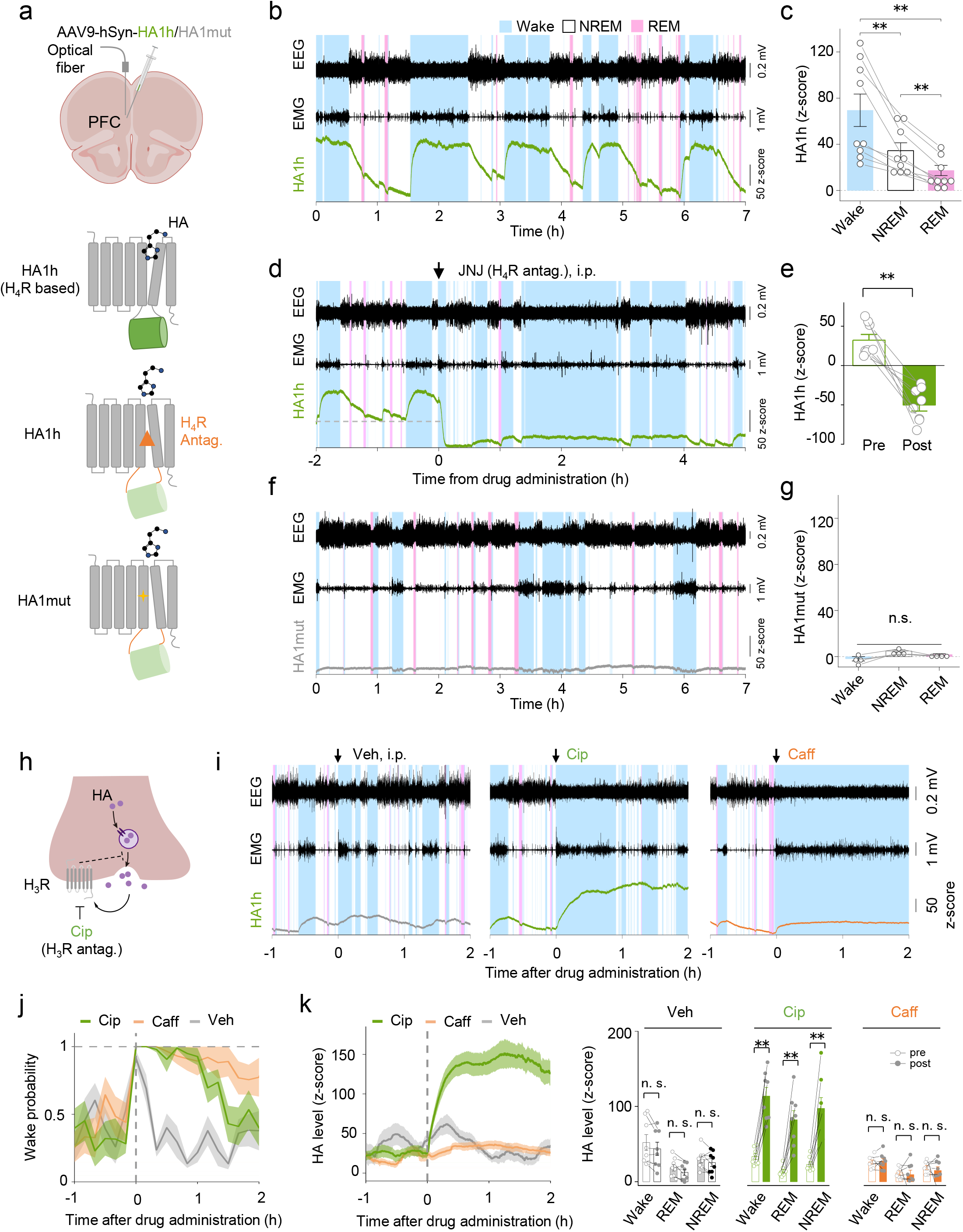
Using the GRAB_HA_ sensor to measure HA release in the PFC in freely moving mice during the sleep-wake cycle. (a) Schematic diagram depicting the strategy for fiber photometry recording of HA1h or HA1mut fluorescence in the prefrontal cortex (PFC) of freely behaving mice during the sleep-wake cycle. Top panel, an AAV expressing either HA1h or HA1mut was injected into the PFC region, and an optical fiber was implanted at the injection site. Bottom three panels, schematic diagram showing the responses the HA1h and HA1mut sensors to HA, as well as the HA1h sensor in the presence of an H_4_R antagonist. (b and f) Example traces of simultaneous EEG, EMG, and HA1h (b) or HA1mut (d) recordings. (c and g) Summary of the average HA1h (c) and HA1mut (e) fluorescence responses measured in the indicated sleep-wake states; (c) *n* = 9 mice. One-way ANOVA: *F* = 21.7374, *P* = 0.0014; Tukey’s *post hoc* test: Wake vs NREM *P* = 0.0074, Wake vs REM *P* = 0.0033, NREM vs REM *P* = 0.0011; ** *P* < 0.01. (e) *n* = 4 mice. One-way ANOVA: *F* = 9.5591, *P* = 0.0512; n.s., not significant. (d) Example traces of simultaneous EEG, EMG, and HA1h recordings after an i.p. injection of the H_4_R antagonist JNJ-7777120 (3 mg/kg body weight). (e) Summary of average HA1h fluorescence measured before (Pre) and after (Post) an injection of JNJ-7777120. *n* = 8 mice Paired t-test, *t* = 6.3859, *P* = 0.0004; ** *P* < 0.01. (h) Schematic drawing depicting the effect of the H_3_R antagonist ciproxifan on downstream HA release. (i) Example traces of simultaneous EEG, EMG, and HA1h recordings in mice before and after administration of vehicle, ciproxifan (3 mg/kg body weight), or caffeine (15 mg/kg body weight). (j) Time course of the probability of being in the awake state; the dashed line at time 0 indicates the administration of vehicle, ciproxifan, or caffeine. (k) Time course (left) and average HA1h signals (right) measured in the PFC before and after administration of vehicle, ciproxifan, or caffeine. *n* = 8. Two-way ANOVA between brain state and time; Vehicle, *F*(time) = 3.9317, *P* = 0.0878; Ciproxifan, *F*(time) = 57.1062, *P* = 0.0001, pre-post comparisons followed by Sidak’s test, Wake, NREM, and REM pre vs post *P* < 0.0001; caffeine, *F*(time) = 0.1526, *P* = 0.7076; ** *P* < 0.01. n.s., not significant. Data are shown as mean ± s.e.m. in c,e,g,j,k with the error bars or shaded regions indicating the s.e.m.

HA is believed to play an important role in both the initiation and maintenance of wakefulness^23^. Thus, the histamine H3 receptor H_3_R is a novel promising target for improving vigilance, and the H_3_R antagonist ciproxifan may have potential therapeutic benefits in the context of vigilance deficiency^28^. In addition, the stimulant caffeine is widely used to maintain wakefulness. We therefore used our HA sensors to determine whether ciproxifan and/or caffeine mediate their effects on wakefulness by increasing HA release (Fig. 5h). As expected, we found that both ciproxifan and caffeine prolonged wakefulness in mice (Fig. 5i,j). Interestingly, however, we found that ciproxifan—but not caffeine—increased the fluorescence signal of HA1h expressed in the PFC (Fig. 5i-k, Supplementary Fig. 6a-c). we further confirm that ciproxifan also increased the fluorescence signal of HA1m expressed in POA in wild type mice, but not HDC KO mice (Supplementary Fig. 6d-f). These results indicate that although both ciproxifan and caffeine promote wakefulness, they do so via distinct mechanisms, with ciproxifan, but not caffeine, increasing HA release in the PFC.

### The kinetics of HA release differ in different brain regions

Histaminergic neurons send their projections to numerous target regions throughout the brain^24^. We therefore asked whether the temporal pattern of HA release is universal throughout the brain or differs among various brain regions. As a first step toward addressing this question, we expressed the HA1h sensor in both the PFC and POA and then simultaneously recorded the fluorescence signals in these two nuclei (Fig. 6a). We found that the magnitude of the HA1h signal measured at each state in the sleep-wake cycle was similar between the PFC and POA (Fig. 6b,c). We also found a close temporal cross-correlation in the time-shift data (Fig. 6d), indicating that the signal change in the POA preceded the signal change in the PFC. We then aligned and normalized the signals measured during the wake-sleep state transitions and found that the signals measured in the POA were faster than the corresponding signals measured in the PFC during both the REM→wake transition, the NREM→wake transition, and the wake→NREM transition (Fig. 6e). We also quantified the kinetics of the responses by calculating the t50 value of the signal rise or fall at each transition. We found that that the t50 values measured in the POA were significantly lower than the corresponding values measured in the PFC during the REM→wake, NREM→wake, and wake→NREM transitions, but were similar during the NREM→REM transition (Fig. 6f), indicating a clear difference in HA release kinetics between the POA and PFC (summarized schematically in Fig. 6g). These results raise at least two possibilities: first, histaminergic projections in the POA and PFC may arise from different subpopulations of histaminergic neurons; alternatively, HA release may be modulated by other neurotransmitters and modulators acting locally at the site of histaminergic innervation (Supplementary Fig. 8).

**Fig. 6:**
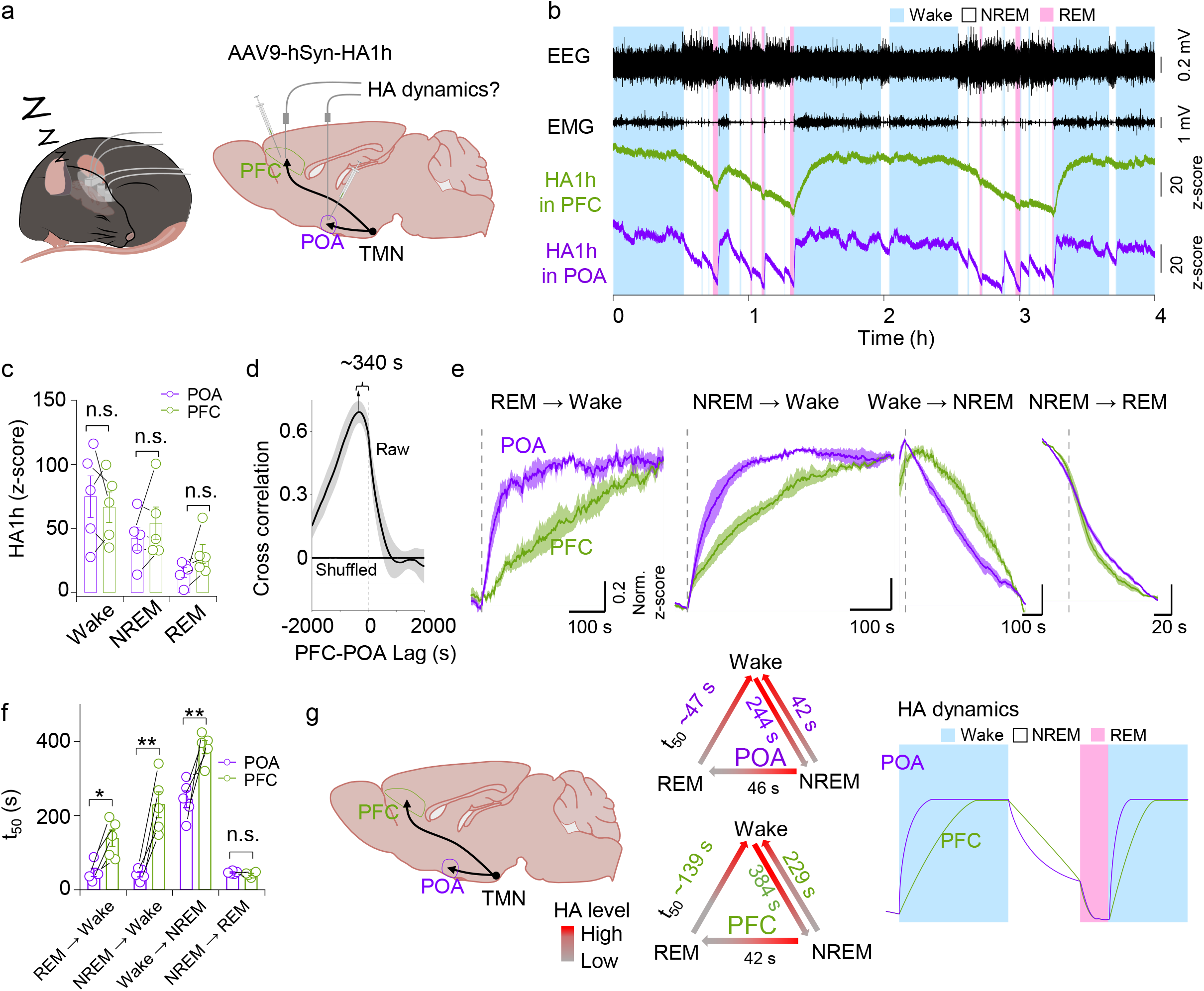
Simultaneously recording HA release in the mouse PFC and POA during the sleep-wake cycle reveals differences in kinetics. (a) Schematic diagram depicting the strategy for simultaneously recording HA1h fluorescence in the PFC and POA during the sleep-wake cycle in a freely moving mouse. (b) Example traces of simultaneous EEG, EMG, and HA1h recordings in the PFC and POA. (c) Summary of average HA1h fluorescence measured in the in the POA and PFC during the indicated sleep-wake states; *n* = 5 mice. Two-way ANOVA between brain state and nuclei, *F*(brain state) = 20.4065, *P* = 0.0007; *F*(nuclei) = 0.4091, *P* = 0.5572. n.s., not significant. (d) Cross-correlation between the HA1h signals measured in the POA and PFC; as a control, the data were also shuffled and depicted as the horizontal line at *y*=0. n = 5 mice. (e) Time courses of the change in HA1h fluorescence measured in the POA and PFC at the indicated transitions between sleep-wake states. The vertical dashed lines represent the transition time. (f) Summary of t50 measured for the change in HA1h fluorescence in the POA and PFC during the indicated sleep-wake state transitions. *n* = 5 mice. Two-way ANOVA between brain state transition and nuclei followed by Sidak’s test, *F*(nuclei) = 52.4873, *P* = 0.0020. REM→Wake, *P* = 0.0156; NREM→Wake, *P* < 0.0001; Wake→NREM, *P* = 0.0006; NREM→REM, *P* = 0.996; * *P* < 0.05, ** *P* < 0.01. (g) Summary of the dynamics measured in the POA and PFC at the transitions between the various sleep-wake states. Data are shown as mean ± s.e.m. in c,d,e,f with the error bars or shaded regions indicating the s.e.m.

## Discussion

Here, we report the development and characterization of two genetically encoded GPCR activation–based fluorescent sensors for measuring changes in extracellular HA in real time. With their high sensitivity, high selectivity, and rapid kinetics, GRAB_HA_ sensors can reliably detect endogenous HA release both *in vitro* and in freely behaving animals under physiological conditions.

Our GRAB_HA_ sensors offer several clear advantages over existing methods for measuring HA. First, GRAB_HA_ sensors have high sensitivity and a relatively high signal-to-noise ratio in response to HA. Specifically, the HA1h and HA1m sensors produce a peak fluorescence response of ~300% and 500%, respectively, with an apparent affinity of ~20 nM and 400 nM, respectively. In contrast, GPCR-based FRET^12^ and BRET^13^ probes for detecting HA have a low peak response of ~160% (or only 25% under optimal conditions) and are limited to use in *in vitro* applications. Second, our GRAB_HA_ sensors have rapid kinetics, with rise and decays times of 0.3-0.6 s and 1.4-2.3 s, respectively. Although this response time of GRAB_HA_ sensors is slightly longer than fast-scan cyclic voltammetry, it is sufficient to report physiologically relevant changes in HA levels and is similar to the response times reported for wild-type GPCRs^29^ and other GRAB sensors^16,20,30^. In addition, our GRAB_HA_ sensors have high spatial resolution and are photostable, allowing for stable recordings over prolonged experiments. Finally, our sensors have high specificity for HA, as they can reliably detect changes in extracellular HA in wild-type mice, but produce no signal in knockout mice lacking the enzyme histidine decarboxylase (HDC). Building on these key advantages over other tools, we used our GRAB_HA_ sensors to measure the release and apparent diffusion rate of HA in acute brain slices by combining the sensor with two-photon imaging. In addition, we combined our sensors with fiber photometry recording to monitor real-time changes in histaminergic activity in the mouse brain during the sleep-wake cycle.

Our GRAB_HA_ sensors have the same high specificity for HA and pharmacological profile of their corresponding parent receptors. Using cultured neurons, we found that both the human and tardigrade receptor-derived HA sensors respond to HA, but do not respond to an HA precursor, HA metabolites, or any other neurotransmitters tested. Similarly, each sensor’s signal was blocked by the corresponding receptor antagonists, but was unaffected by other HA receptor antagonists, indicating that the complementary pharmacological profile of these two HA sensors can be exploited in order to study how HA release is regulated by various compounds that target histaminergic signaling. For example, the H_3_R receptor is a promising new target for improving vigilance, and an H_3_R antagonist has been shown to increase HA turnover by blocking the negative feedback regulation of HA release (see Fig. 5h)^31,32^. Consistent with these previous studies, we found that the GRAB_HA_ response was larger in wild-type mice that received an H_3_R antagonist, but was not affected in HDC-deficient mice (see Supplementary Fig. 6d-f), indicating that our HA sensors can be used to monitor the effects of H_3_R antagonists on histaminergic signaling.

Using our GRAB_HA_ sensor, we also found that specific brain regions have distinct patterns of HA release kinetics. This finding suggests that the projections of histaminergic neurons are heterogenous; however, an alternative possibility is that HA release may be modulated to some extent by local or regional neurotransmitters, modulators, and/or pathways (Supplementary Fig. 8). In the brain, histaminergic projections: *i*) originate from a single source, namely the tuberomammillary nucleus (TMN) in the posterior hypothalamus; *ii*) innervate nearly all regions of the brain; and *iii*) regulate a wide range of behaviors and processes. Histaminergic TMN neurons are organized into five clusters, E1 through E5 ^33,34^; however, retrograde tracers injected into various brain regions labeled the entire TMN, without any clear topographical pattern^33,35^. By combining cell type–specific sparse labeling with fluorescence micro-optical sectioning tomography, whole-brain histaminergic axonal projection mapping revealed that histaminergic neurons indeed send heterogeneous projections to their downstream target regions^36^. With respect to the second possibility, several studies have shown that neurotransmitter release can be modulated at the level of the axon terminal. For example, dopamine release in the striatum is regulated at the axon terminal by acetylcholine^37,38^. Thus, HA release may also be modulated by the local environment, giving rise to our observation of heterogeneous release dynamics among target regions. Future studies combining our GRAB_HA_ sensors with other tools and strategies may shed new light on the mechanism underlying HA release and its regulation at the local and/or regional level.

In addition to regulating the sleep-wake cycle, HA plays a role in a number of physiological and pathological processes in the central nervous system, including feeding, motor control, cognition, narcolepsy, epilepsy, migraine, and Tourette’s syndrome^39,40^. Interestingly, HA also serves as the primary neurotransmitter for visual input in *Drosophila* photoreceptors^41,42^. Moreover, histamine plays a major role in allergic responses, including anaphylaxis^1^. Thus, this novel set of HA sensors suitable for both *in vitro* and *in vivo* applications can be used to measure HA release in a variety of settings and models, providing valuable new insights into the role of histaminergic signaling in both health and disease.

## Methods

### Animals

All experimental procedures involving animals were performed in accordance with the guidelines established by the Animal Care and Use Committee at Peking University. Wild-type C57BL/6j mice were obtained from the Beijing Vital River Laboratory and used to prepare the acute brain slices and for some *in vivo* mouse experiments. Histidine decarboxylase knockout (HDC KO) mice (on a BALB/c background) were kindly provided by Dr. Xiangdong Yang at Fudan University; HDC KO and control littermates were used for *in vivo* recordings. Newborn wild-type Sprague-Dawley rat pups (P0) were obtained from the Beijing Vital River Laboratory and used to prepare primary cortical neuron cultures. All animals were housed at 18–23°C in 40–60% humidity under a 12/12-h light/dark cycle, with food and water available *ad libitum*.

### Molecular biology

The clones used in this study were generated using the Gibson assembly method. DNA fragments were amplified by PCR using primers (TSINGKE Biological Technology) with 25–30-bp overlap and then assembled using T5 exonuclease (New England Biolabs), Phusion DNA polymerase (Thermo Fisher Scientific), and Taq ligase (iCloning). All plasmid sequences were verified using Sanger sequencing (TSINGKE Biological Technology). To characterize the sensors in HEK293T cells, all cDNAs encoding the candidate GRAB_HA_ sensors were cloned into the pDisplay vector with an upstream IgK leader sequence and a downstream IRES-mCherry-CAAX cassette to label the cell membrane and calibrate the sensor’s fluorescence intensity. To characterize the sensors in cultured neurons, sequences encoding HA1h, HA1m, and HA1mut were cloned into the pAAV vector containing the human synapsin (hSyn) promoter. To measure downstream signaling using the Tango assay, genes encoding wild-type H_1_R and H_4_R, as well as the HA1h and HA1m sensors, were cloned into the pTango vector. To characterize the sensors’ wavelength spectra, sequences encoding HA1h and HA1m were cloned into the pPacific vector containing a 3’ terminal repeat, IRES, puromycin gene, and a 5’ terminal repeat. Two mutations (S103P and S509G) were introduced into the pCS7-PiggyBAC vector (ViewSolid Biotech) to generate a hyperactive piggyBac transposase, which was used to generate stable cell lines expressing HA1h or HA1m.

### Recombinant adeno-associated virus (AAV)

AAV2/9-hSyn-HA1h (2.64×10^13^ GC/ml, WZ Biosciences), AAV2/9-hSyn-HA1m (3.92×10^13^ GC/ml, WZ Biosciences), and AAV2/9-hSyn-HA1mut (9.03×10^13^ GC/ml, WZ Biosciences) were used to infect cultured neurons or were injected into specific mouse brain regions.

### Cell culture

HEK293T cells were purchased from ATCC and verified based on their morphology observed by microscopy and an analysis of their growth curve; the cells were cultured at 37°C in humidified air containing 5% CO_2_ in DMEM (Biological Industries) supplemented with 10% (v/v) FBS (Gibco), penicillin (100 U/ml), and streptomycin (0.1 mg/ml) (Biological Industries). For expressing the GRAB_HA_ sensors, HEK293T cells were plated on 96-well plates or 12-mm glass coverslips in 24-well plates and grown to 60-70% confluence, and then transfected using polyethylenimine (PEI) with 300 ng DNA per well (for 96-well plates) or 1 μg DNA per well (for 24-well plates) at a DNA:PEI ratio of 1:3. The culture medium was replaced with fresh medium 6-8 h after transfection, and fluorescence imaging was performed 24–48 h after transfection.

Rat cortical neurons were prepared from P0 Sprague-Dawley rat pups. In brief, the cortex was dissected, and the neurons were dissociated in 0.25% trypsin–EDTA (Gibco), plated on 12-mm glass coverslips coated with poly-D-lysine (Sigma-Aldrich), and cultured in neurobasal medium (Gibco) containing 2% B-27 supplement (Gibco), 1% GlutaMAX (Gibco), and 1% penicillin-streptomycin (Gibco) at 37°C in humidified air containing 5% CO_2_. For viral infection, cultured neurons were infected with AAV2/9-hSyn-HA1h, AAV2/9-hSyn-HA1m, or AAV2/9-hSyn-HA1mut at 3–5 days *in vitro* (DIV3–DIV5), and fluorescence imaging was performed at DIV10–DIV14.

### Fluorescence imaging of cultured cells

Cultured cells were imaged using the Opera Phenix high-content screening system (PerkinElmer) and an inverted Ti-E A1 confocal microscope equipped with a 20x 0.4-numerical aperture (NA) objective, a 40x 0.6-NA objective, a 40x 1.15-NA water-immersion objective, a 488-nm laser, and a 561-nm laser. A 525/50 nm emission filter and a 600/30 nm emission filter were used to collect the GFP and RFP signals, respectively.

During fluorescence imaging, cells expressing the GRAB_HA_ sensors were first bathed in Tyrode’s solution and then imaged before and after addition of the indicated drugs at various concentrations. The change in fluorescence intensity of the GRAB_HA_ sensors was calculated using the change in the GFP/RFP ratio and is expressed as ΔF/F_0_.

To measure the response kinetics of GRAB_HA_ sensors in the rapid perfusion experiments using the line-scanning mode, the tip of a glass pipette containing 100 μM HA was placed near the sensor-expressing HEK293T cells. HA was puffed onto the cells from the pipette to measure the on-rate. To measure the off-rate, 1 mM of the appropriate antagonist (JNJ-7777120 to block HA1h, clemastine to block HA1m) was applied using a glass pipette to sensor-expressing cells bathed in HA (1 μM HA for cells expressing HA1h, 10 μM HA for cells expressing HA1m).

### Spectra measurements

For the HA1h sensor, HEK293T cells stably expressing HA1h under the control of CAG promoter were plated in a 384-well plate in the absence or presence of 100 μM HA. Control cells used for background subtraction were transfected with an empty vector. Using a Safire^2^ multi-mode plate reader (Tecan), excitation spectra were measured from 300 to 520 nm at 5-nm increments and a bandwidth of 20 nm, while the emission spectrum was set to 560 nm with a 20-nm bandwidth. Emission spectra were measured from 500 to 700 nm at 5-nm increments and a bandwidth of 20 nm, while the excitation spectrum was set to 455 nm with a bandwidth of 20 nm. For the HA1m sensor, the plasmid expressing HA1m or an empty vector was transfected into HEK293T cells in 6-well plates; 24–36 h after transfection, the cells were harvested using trypsin, washed with PBS, resuspended in Tyrode’s solution in the absence or presence of 100 μM HA, and plated into a 384-well plate, followed by the spectra measurements described above.

### G_qi_ calcium imaging assay

Plasmids expressing the GRAB_HA_ sensors, H_1_R-EGFP, or H_4_R-EGFP were co-transfected with a construct expressing jRGECO1a-P2A-G_q-i1_ into HEK293T cells plated on 12-mm coverslips; 24–48 h after transfection, the cells were imaged using a Ti-E A1 confocal microscope (Nikon) as described above. The indicated concentrations of HA were then applied by bath application and removed using a custom-made perfusion system.

### Tango assay

Plasmids expressing wild-type H_1_R, wild-type H_4_R, HA1h, or HA1m were transfected into HTLA cells in 6-well plates; 24 h after transfection, the cells were harvested using trypsin and plated in 96-well plates. HA was then applied at a final concentration ranging from 1 nM to 1 mM, and the cells were cultured for an additional 12 h to allow for luciferase expression. Bright-Glo (Luciferase Assay System, Promega) was then added to a final concentration of 5 μM, and luminescence was measured using a VICTOR X5 multilabel plate reader (PerkinElmer).

### [^3^H]histamine competition binding assay

Plasmids expressing wild-type H_4_R or HA1h were transfected into HEK293T cells in 10 cm dishes; 48 h after transfection, the cells were collected, centrifuged at 2000*g* for 10 min at 4°C, resuspended in 50 mM Tris-HCl binding buffer (pH7.4), and sonicated for 10 sec using a Branson sonifier 250 (Boom bv., Meppel, The Netherlands). The cell homogenates were incubated with ~8.5 nM [^3^H]-histamine in the absence of presence of 30 μM unlabeled compounds in a total volume of 100 μl/well at 25°C on a shaker (600 rpm). After 2 h, the incubations were terminated by rapid filtration through 0.5% (v/v) polyethylenimine pre-soaked GF/C filterplates using a 96-well Filtermate harvester (PerkinElmer), followed by four rapid washes with ice-cold binding buffer. The GF/C filterplates were dried at 52°C and 25 μl Microscint-O was added to each well to quantify radioactivity using a Microbeta Wallac Trilux scintillation counter (PerkinElmer). Saturation binding of [^3^H]histamine was measured with increasing concentrations of [^3^H]histamine in the absence or presence of 50 microM JNJ7777120 to detect total and nonspecific binding respectively. The binding affinity (K_d_) values were determined using nonlinear fitting (One site – total and nonspecific binding) in GraphPad Prism 9.4.0.

### Two-photon fluorescence imaging of mouse acute brain slices

For AAV injection, adult wild-type C57BL/6N mice were firstly anesthetized by an i.p. injection of Avertin (500 mg/kg, Sigma-Aldrich), and then fixed in a stereotaxic frame (RWD Life Science) for injection of the AAV expressing hSyn-HA1m (300 nl per site) using a micro-syringe pump (Nanoliter 2000 Injector, World Precision Instruments). The AAV was injected into the PFC of the left hemisphere of C57BL/6N mice using the following coordinates: AP: +1.9 mm relative to Bregma, ML: −0.3 mm, and DV: −1.8 mm below the dura.

For slice preparation, 2-3 weeks after virus injection, mice were deeply anesthetized again with an i.p. injection of Avertin, and then transcardial perfusion was rapidly performed using cold oxygenated slicing buffer containing (in mM): 110 Choline-Cl, 25 NaHCO_3_, 25 Glucose, 7 MgCl_2_, 2.5 KCl, 1.3 Na ascorbate, 1 NaH_2_PO_4_, 0.5 CaCl_2_, and 0.6 Na pyruvate. The isolated brains were subsequently immersed into the oxygenated slicing buffer, and the cerebellum was trimmed using a razor blade. The caudal sides were then glued on the cutting stage of a VT1200 vibratome (Leica) and sectioned into 300-μm thick coronal slices. Brain slices containing the PFC region were incubated at 34°C for ~40 min in the oxygen-saturated Ringer’s buffer containing (in mM): 125 NaCl, 25 NaHCO_3_, 25 Glucose, 2.5 KCl, 2 CaCl_2_, 1.3 MgCl_2_, 1.3 Na ascorbate, 1 NaH_2_PO_4_, and 0.6 Na pyruvate.

For two-photon imaging, the HA1m-expressing slices were transferred into a perfusion chamber in an Ultima Investigator 2P microscope (Bruker) equipped with a 25× /1.05-NA water-immersion objective and a mode-locked Mai Tai Ti: Sapphire laser (Spectra-Physics). The laser wavelength was tuned to 920 nm for measuring the fluorescence of HA1m using a 495-540 nm filter. A homemade bipolar electrode (cat. #WE30031.0A3, MicroProbes) was placed onto the slice surface near the PFC under fluorescence guidance. Fluorescence was then recorded at video frame rates of 0.3259 s/frame, with a resolution of 256×256 pixels. The stimulation voltage was set at 4-6 V with the duration of each stimulation at 1 ms. Stimulation and imaging were synchronized through a custom-written program running on an Arduino board.

### Surgery for virus injection, placement of EEG and EMG electrodes, and fiber implantation

Adult mice were anesthetized with 1.5% isoflurane, and 2% lidocaine hydrochloride was injected subcutaneously under the scalp; the mouse was then placed on a stereotaxic frame (RWD Life Science). The skull was exposed, and small craniotomy holes were prepared for virus injection, into which a total of approximately 300 nl AAV9-hSyn-HA1h, AAV9-hSyn-HA1m, or AAV9-hSyn-HA1mut virus was microinjected into the PFC (AP: +1.9 mm relative to Bregma, ML: −0.3 mm, DV: 1.9 mm below the dura) and/or POA (AP: 0 mm relative to Bregma, ML: −0.6 mm, DV: 4.9 mm below the dura) via a fine glass pipette and micro-syringe pump (Nanoliter 2010 injector, World Precision Instruments).

For recording the fluorescence signals, a 200-μm optical fiber cannula (Fiber core: 200 μm; numerical aperture: 0.37; Inper, Zhejiang, China) was implanted 0.1 mm above the virus injection sites.

To record the animal’s sleep-wake state, EEG electrodes were implanted into the craniotomy holes over the frontal cortex and visual cortex, and EMG wires were placed bilaterally into the trapezius muscles. The electrodes were attached to a microconnector and fixed to the skull using dental cement.

### Fiber photometry recording and analysis

A fiber photometry system (Thinker Tech, Nanjing, China) was used to record the fluorescence signals in freely moving mice. Blue (473-nm) LED light (Cree LED) was bandpass filtered (470/25 nm, model 65-144, Edmund Optics), reflected by a dichroic mirror (model 67-069, Edmund Optics), and then focused using a 20x objective lens (Olympus). An optical fiber guided the light between the commutator and the implanted optical fiber cannula. The excitation light power at the tip of the optical fiber was adjusted 20-30 μW and delivered at 100 Hz with 5 ms pulse duration when the recording lasted <24 h; the light power was adjusted to a lower level (10-20 μW) and was delivered at 10 Hz when the recording lasted >24 h in order to minimize photobleaching. The green fluorescence was bandpass filtered (525/25 nm, model 86-354, Edmund Optics) and collected using a photomultiplier tube (model H10721-210, Hamamatsu). An amplifier (model C7319, Hamamatsu) was used to convert the current output from the photomultiplier tube to a voltage signal and was passed through a low-pass filter. The analog voltage signals were then digitized using an acquisition card (National Instruments). To minimize autofluorescence of the optical fiber, the recording fiber was photobleached using a high-power LED before recording. Background autofluorescence was subtracted from the recorded signals in the subsequent analysis.

The photometry data were analyzed using a custom program written in MATLAB. To calculate ΔF/F_0_, baseline values were measured during REM sleep with no apparent fluctuations. To compare the change in fluorescence between animals, the *z*-score–transformed ΔF/F_0_ was further normalized using the standard deviation of the baseline signals.

### Polysomnographic recording and analysis

We used the EEG and EMG recordings to determine the animal’s sleep-wake state. The EEG and EMG electrode microconnectors were connected via a flexible cable and attached to an electric slip ring to allow the mice to move freely. The cortical EEG and neck EMG signals were amplified (NL104A, Digitimer), filtered (NL125/6, Digitimer), digitized using a Power1401 digitizer (Cambridge Electronic Design Ltd.), and recorded using Spike2 software (Cambridge Electronic Design Ltd.) at a sampling rate of 1000 Hz. The polysomnographic signals were filtered (EEG: 0.5-100 Hz, EMG: 30-500 Hz) and semi-automatically scored off-line in 4-s epochs of wakefulness, REM sleep, and NREM sleep using AccuSleep (https://github.com/zekebarger/AccuSleep) ^43^; the defined sleep-wake stages were examined visually and corrected if necessary. Wakefulness was defined as desynchronized low-amplitude EEG activity and high-amplitude EMG activity with phasic bursts. NREM sleep was defined as synchronized EEG with high-amplitude delta rhythm (0.5-4 Hz) and low EMG activity. REM sleep was defined as a pronounced theta rhythm (6-10 Hz) and low EMG activity. EEG spectral analysis was estimated using a short-time fast Fourier transform (FFT).

### Quantification and statistical analysis

The animals and cells in the experiments were randomly assigned to different groups at sample sizes that were determined based on previous studies^44^. Imaging data from cultured HEK293T cells, cultured rat cortical neurons, and acute mouse brain slices were processed using ImageJ software (NIH) and analyzed using custom MATLAB code. Exponential-function fitting was used to correct for slight photobleaching. Background levels measured outside the region of interest in the pseudocolor images were subtracted using ImageJ.

Origin 2020 (OriginLab) and Prism 8 & 9 (GraphPad) were used to perform the statistical analyses. Except where indicated otherwise, all summary data are presented as the mean ± sem. The paired or unpaired Student’s *t*-test was used to compare two groups, and a one-way analysis of variance (ANOVA) was used to compare more than two groups. All statistical tests were two-tailed, and differences were considered statistically significant at *P* < 0.05.

## Data and code availability

Plasmids expressing the sensors used in this study were deposited at Addgene (Addgene ID, HA1h, 190473. HA1m, 190474. HA1mut 190475). The data and custom-written ImageJ macro, Arduino, and MATLAB programs are available upon request to the corresponding author.

## Acknowledgments

This research was supported by grants from the National Natural Science Foundation of China (31925017, 31871087, and 81821092); grants from the NIH BRAIN Initiative (1U01NS113358 and 1U01NS120824); grants from National Key R&D Program of China (2020YFE0204000 and 2019YFA0801603) the FENG Foundation; and grants from the Peking-Tsinghua Center for Life Sciences and the State Key Laboratory of Membrane Biology at Peking University School of Life Sciences. We thank Yi Rao for sharing the two-photon microscope, Xiaoguang Lei at PKU-CLS, and the National Center for Protein Sciences at Peking University in Beijing, China, for support and assistance with the Opera Phenix high-content screening system and imaging platform. We thank Xiangdong Yang at Fudan University kindly provided HDC KO mice. Some graphics were generated using BioRender.com.

## Author contributions

Y.L. conceived and supervised the project. M.L. performed the experiments related to developing, optimizing and characterizing the sensor in cultured HEK293T cells and neurons. Y. Y. characterized HA sensors in HEK293T cells and neurons. X.M and H.F.V performed the binding studies. T.Q. and M.L. performed the experiments in brain slices. H.D., Y.L., and C.L. performed the *in vivo* recording in behaving mice. G.L. and H.W. performed the characterization of the wavelength spectra of HA sensors in HEK293T cells. All authors contributed to data interpretation and analysis. Y.L., H.D., and Y.Y. wrote the manuscript with input from all other authors.

---

**Supplementary Fig. 1:**
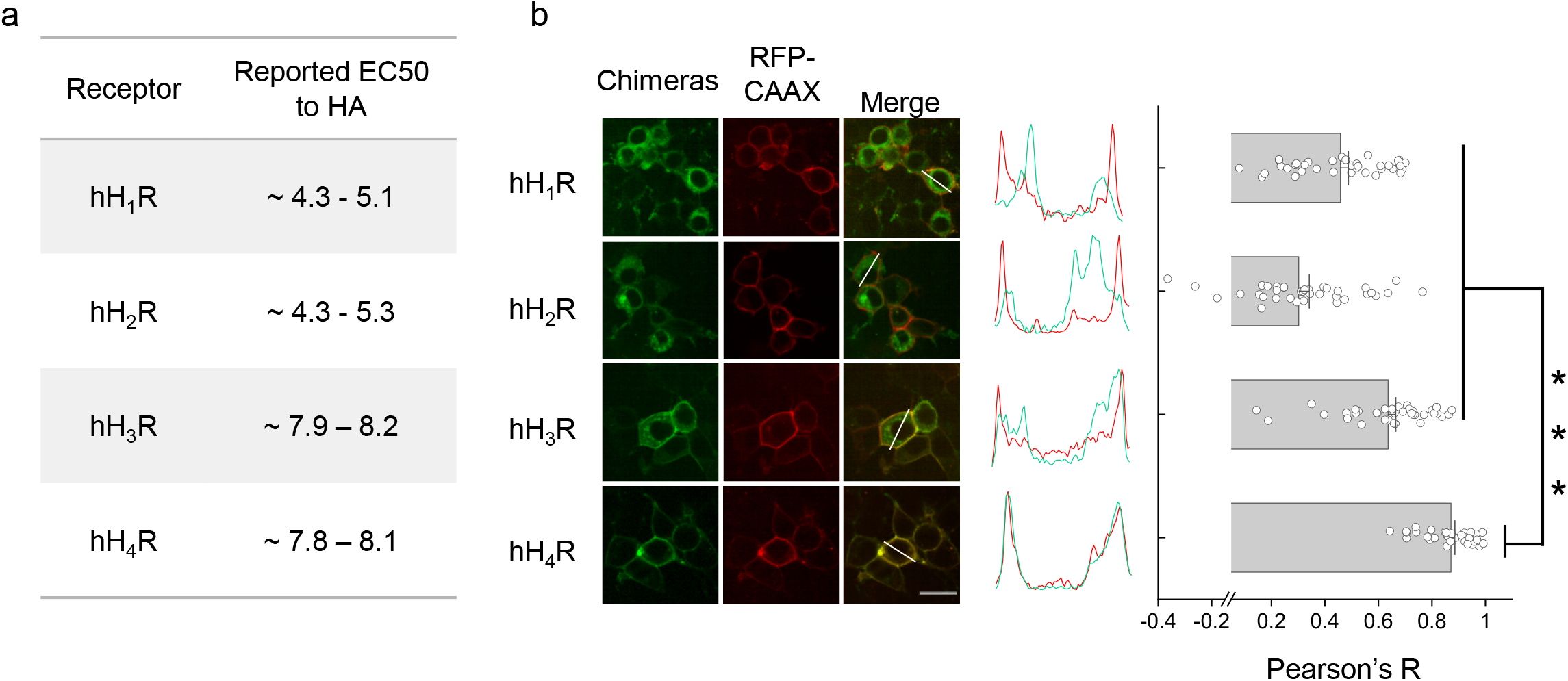
Screening of HA sensors with good membrane trafficking from four human HA receptors. (a) Summary of the reported EC50 values for the indicated human HA receptors (source: https://www.guidetopharmacology.org/). (b) Analysis of fluorescence and membrane trafficking of all four human HA receptors containing cpEGFP expressed in HEK293T cells; membrane-targeted RFP (RFP-CAAX) was co-expressed to label the plasma membrane. Left, fluorescence images of HEK293T cells expressing the indicated HA receptor-based chimeras (green) and RFP (red). Middle, normalized line-scanning plots of the fluorescence signals measured in both the green and red channels. Right, summary of Pearson’s co-localization ratio measured between the indicated HA receptor-based chimeras and RFP-CAAX; *** *P* <0.001 versus hH_4_R. Scale bar, 20 μm.

**Supplementary Fig. 2:**
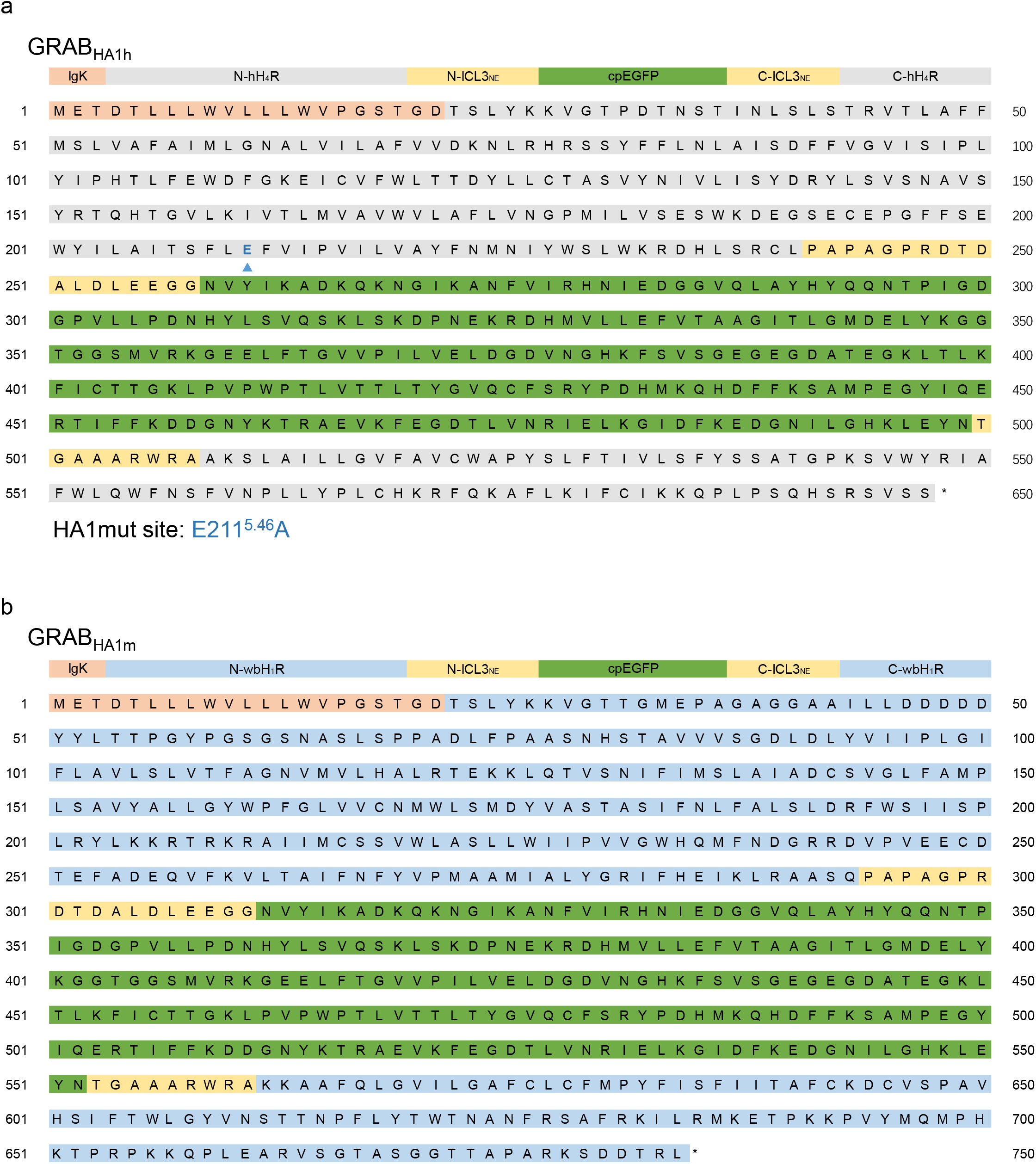
The amino acid sequences of the GRAB_HA_ sensors. The amino acid sequences of the GRAB_HA1h_ (a) and GRAB_HA1m_ (b) sensors are shown, with the indicated domains shown above. Also shown is the location of the E211A mutation (blue arrowhead) in GRAB_HA1h_ to generate the GRAB_HA1mut_ sensor.

**Supplementary Fig. 3:**
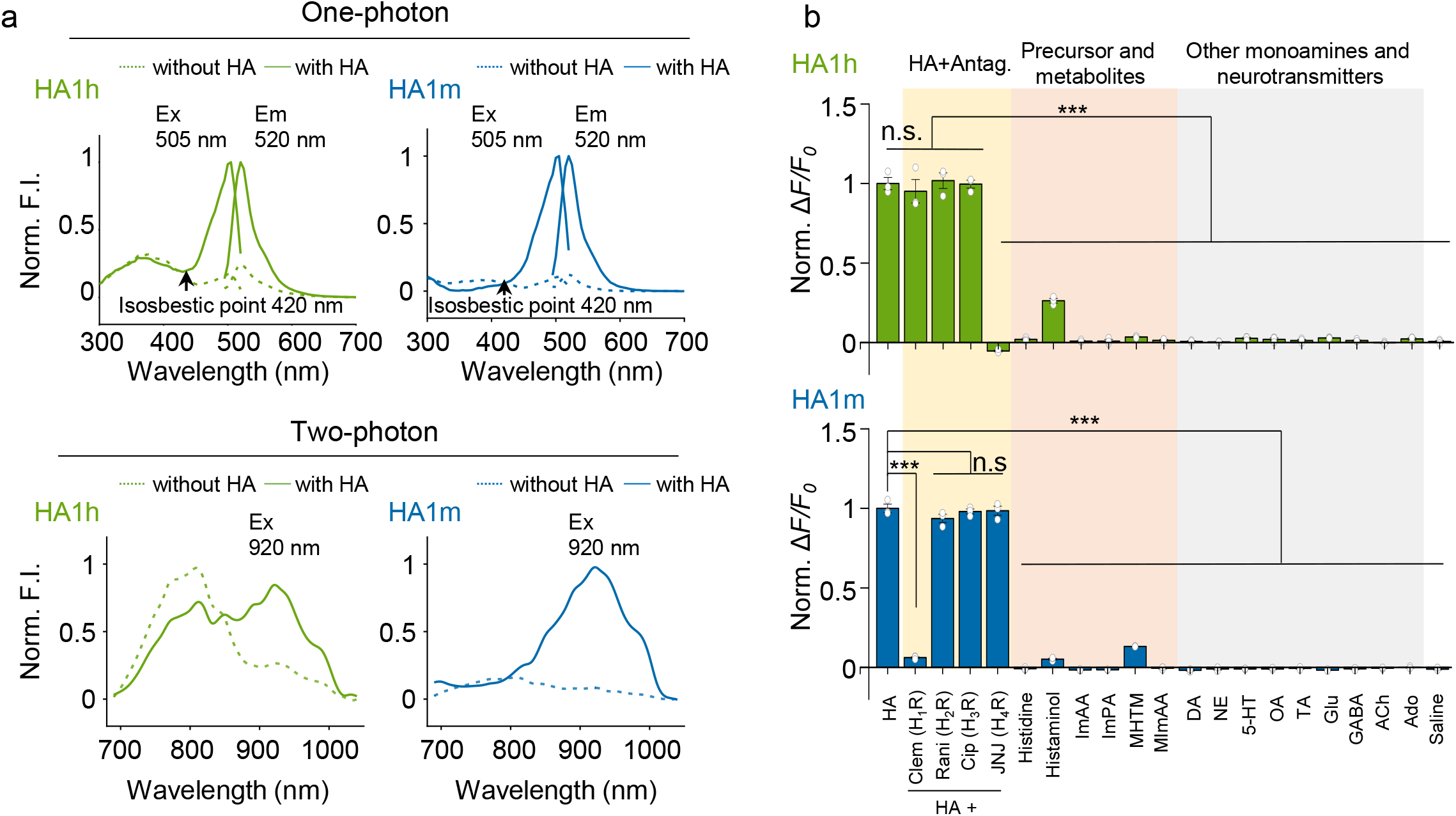
Characterization of the wavelength spectra and specificity of HA sensors expressed in HEK293T cells. (a) One-photon excitation (Ex) and emission (Em) spectra (left) and two-photon excitation spectra (right) of HA1h and HA1m measured in the absence and presence of HA. (b) Summary of the normalized change in fluorescence measured for HA1h (top panel) and HA1m (bottom panel) expressed in HEK293T cells in response to the indicated compounds (applied at 5 μM each); see Fig. 2 for abbreviations. n = 3 wells for different groups. Paired two-tailed Student’s t-tests were performed. *** *P* < 0.001; n.s., not significant.

**Supplementary Fig. 4:**
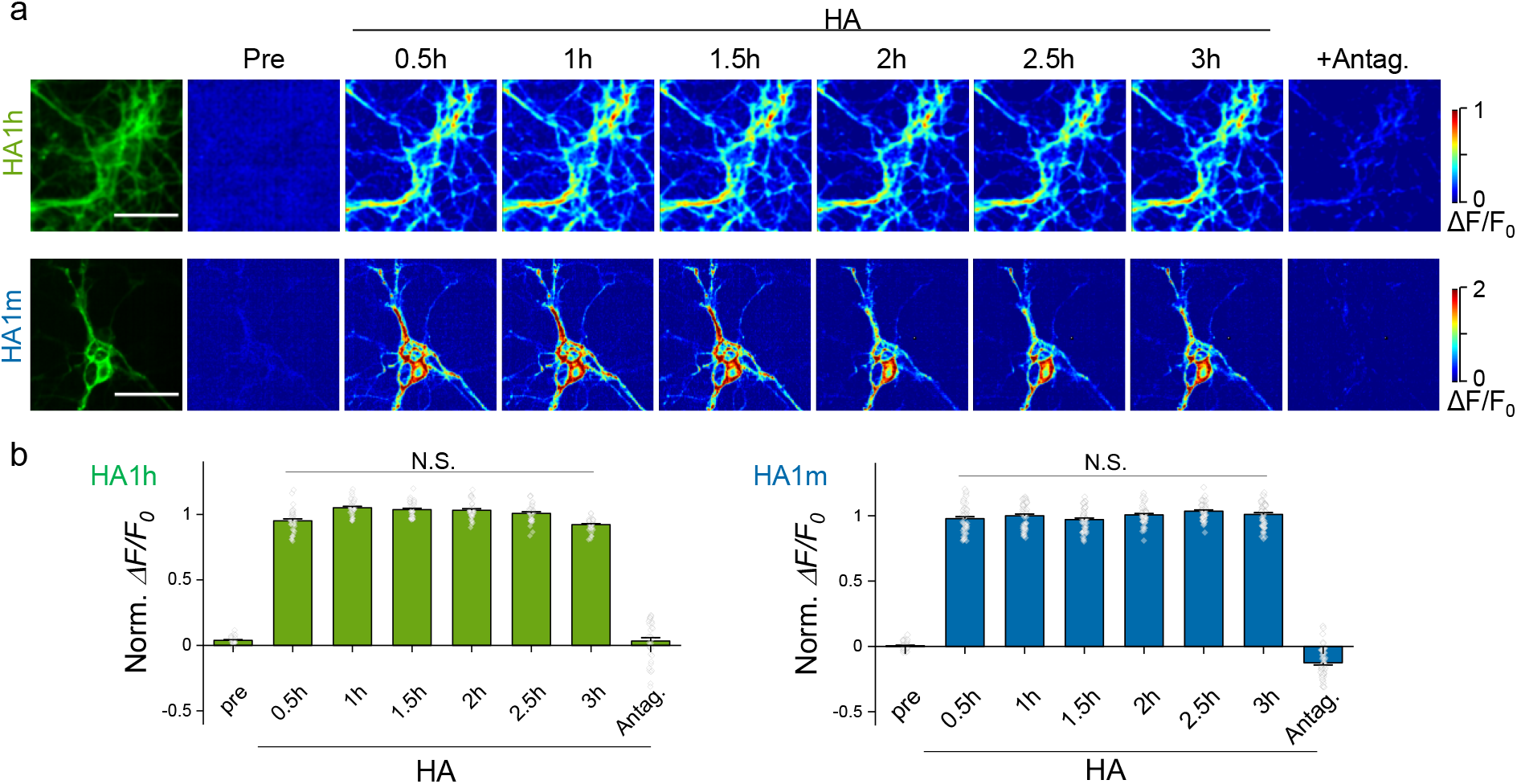
3-h recording of HA1h and HA1m expressed in cultured cortical neurons. Shown are fluorescence images (a) and summary data (b) of the indicated HA sensors expressed in cultured neurons before (Pre) and during a 3-h application of HA at saturating concentration, followed by application of the corresponding antagonist.

**Supplementary Fig. 5:**
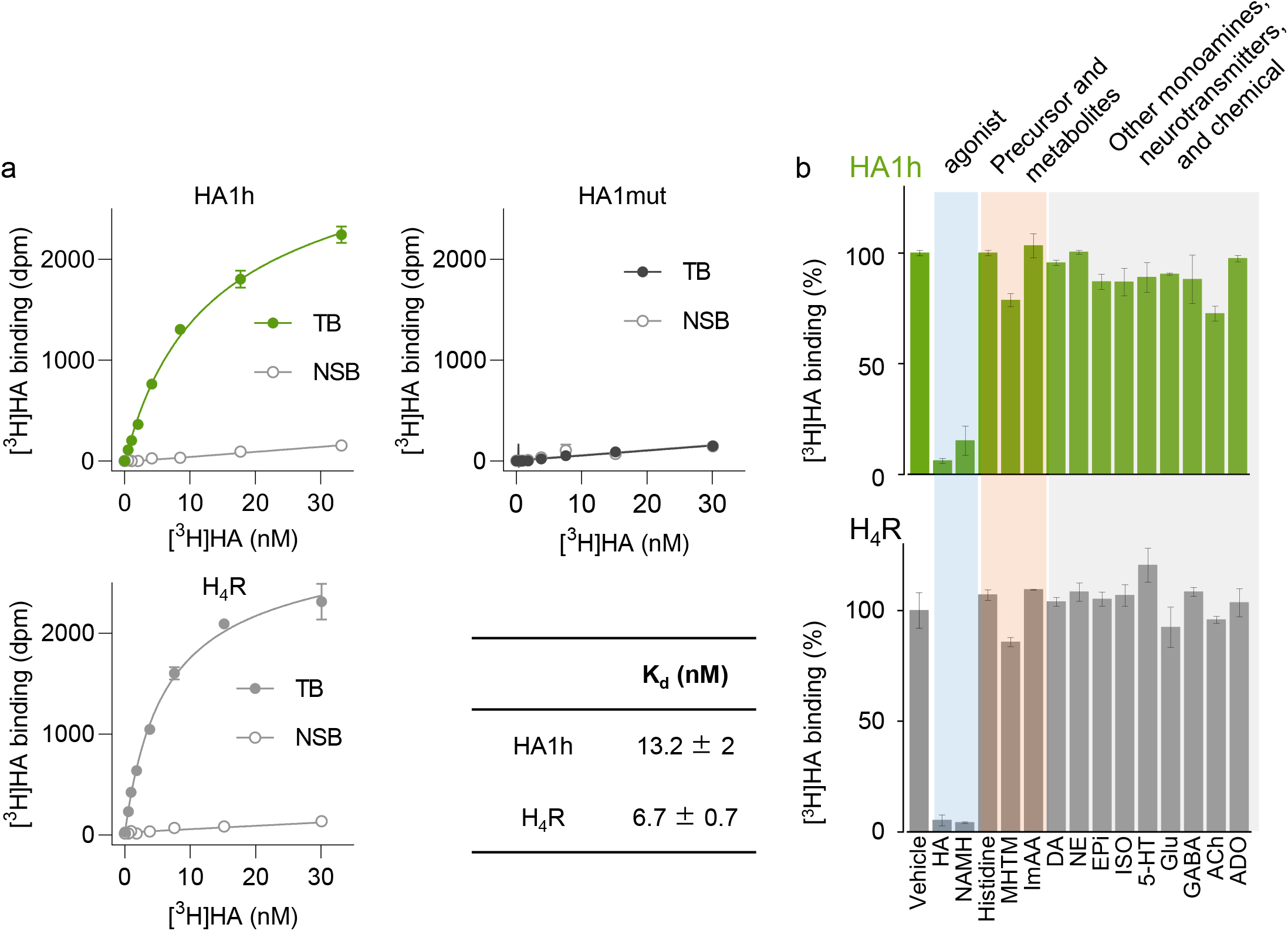
[^3^H]HA radioligand binding assay for HA1h sensor. (a) Binding affinity measurement. Specific binding of increasing concentrations [^3^H]HA to HEK293T cell homogenates expressing HA1h, HA1mut, or H_4_R. TB: total binding, NSB: nonspecific binding. (b) [^3^H]HA radioligand competition binding assay for measurement of [^3^H]HA (8.5 nM) binding to test whether HA, H_4_R agonist, HA precursor, HA metabolites, and other monoamines, neurotransmitters, and chemicals (all compounds were applied at 30 μM) bind to HA1h sensor (top panel) and WT H_4_R (bottom panel).

**Supplementary Fig. 6:**
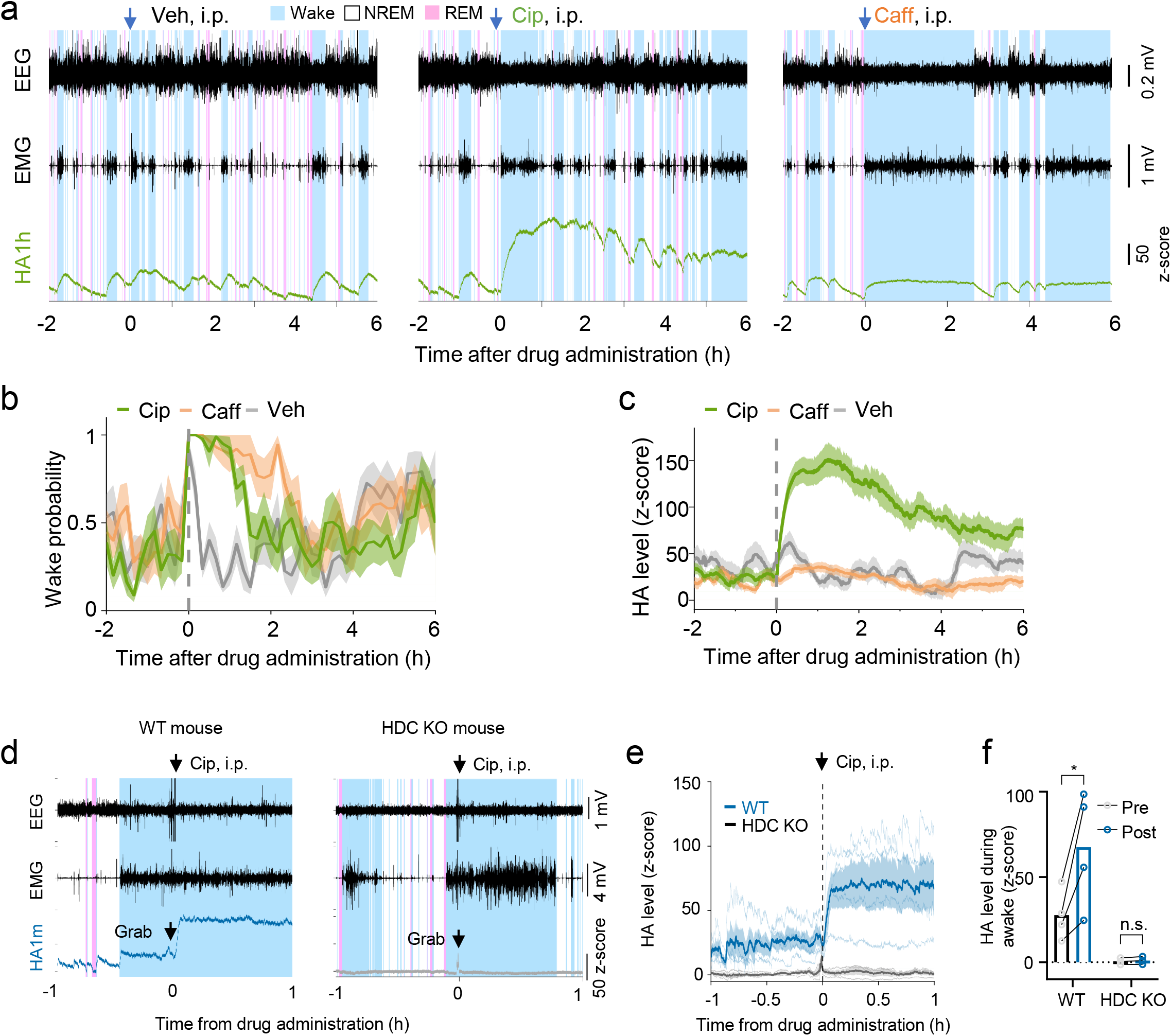
Wake-promoting agent, H_3_R antagonist, enhanced HA level in the PFC of mice, extend shown time in Fig. 5i,j,k. (a) Typical traces of EEG, EMG and HA1h signals after administration of wake-promoting agents, H3R antagonist, ciproxifan at 3 mg/kg, or caffeine at 15 mg/kg, or vehicle in WT mice, extend shown time in Fig. 5i. (b) Time courses of wake probability before and after administration of ciproxifan, caffeine and vehicle, extend shown time in Fig. 5j. (c) Time courses of HA1h signals before and after administration of ciproxifan, caffeine and vehicle, extend shown time in Fig. 5k. (d) Typical traces of EEG, EMG and HA1m signals after administration of H3R antagonist, ciproxifan at 3 mg/kg in WT mouse (left panel) and HDC KO mouse (right panel). (e) Time courses of HA1m signals before and after administration of ciproxifan in WT mouse and HDC KO mouse. (f) Ground data of HA1m signals before and after administration of ciproxifan in WT mouse and HDC KO mouse. Two-way ANOVA between genotype and time; *F*(time) = 9.6457, *P* = 0.0267, pre-post comparisons followed by Sidak’s test WT pre vs post *P* = 0.0107, HDC KO pre vs post *P* = 0.9989;

**Supplementary Fig. 7.**
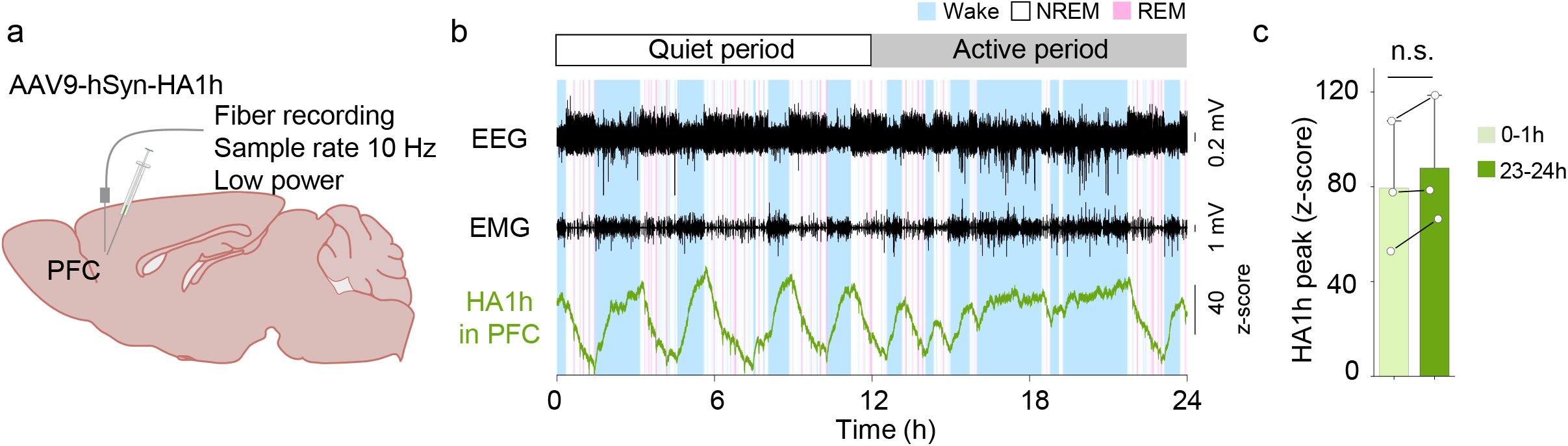
HA1h can be used to measure HA dynamics over one day *in vivo*. (a) Schematic diagram depicting Simultaneous recording of HA1h in the PFC and POA during sleep–wake cycles. (b) Typical traces of EEG, EMG and HA1h signals over 24h. The brain states are color-coded. (c)Summary data of first and last hour averaged HA1h signals. Paired two-tailed Student’s t-tests were performed. *t* = 2.184, *P* = 0.161. n.s., not significant.

**Supplementary Fig. 8:**
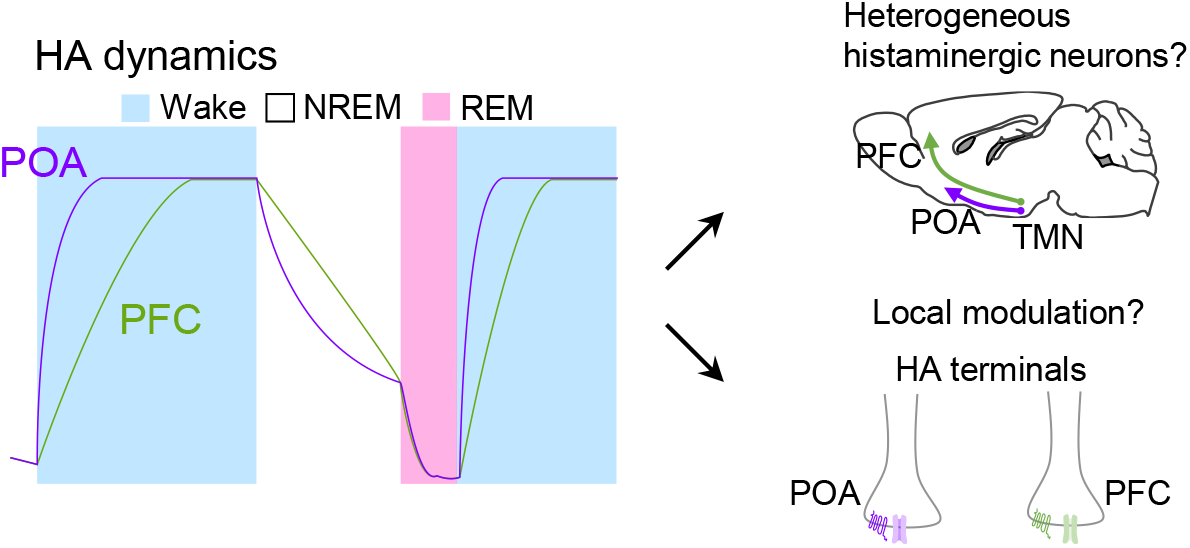
Model depicting two possible explanations for the difference in HA release kinetics between the POA and PFC.

## Reference

1 White, M. The role of histamine in allergic diseases. Journal of Allergy and Clinical Immunology 86, 599–605, doi:10.1016/s0091-6749(05)80223-4 (1990).

2 Prinz, C., Zanner, R. & Gratzl, M. Physiology of gastric enterochromaffin-like cells. Annu Rev Physiol 65, 371–382, doi:10.1146/annurev.physiol.65.092101.142205 (2003).

3 Faingold, C. L. in Histamine II and Anti-Histaminics: Chemistry, Metabolism and Physiological and Pharmacological Actions (ed Mauricio Rocha e Silva) 561–573 (Springer Berlin Heidelberg, 1978).

4 Watanabe, T. et al. Evidence for the presence of a histaminergic neuron system in the rat brain: An immunohistochemical analysis. Neuroscience Letters 39, 249–254, doi:10.1016/0304-3940(83)90308-7 (1983).

5 Haas, H. & Panula, P. The role of histamine and the tuberomamillary nucleus in the nervous system. Nat Rev Neurosci 4, 121–130, doi:10.1038/nrn1034 (2003).

6 Yoshitake, T. et al. Determination of histamine in microdialysis samples from rat brain by microbore column liquid chromatography following intramolecular excimer-forming derivatization with pyrene-labeling reagent. J Neurosci Methods 127, 11–17, doi:10.1016/s0165-0270(03)00097-9 (2003).

7 Puthongkham, P., Lee, S. T. & Venton, B. J. Mechanism of Histamine Oxidation and Electropolymerization at Carbon Electrodes. Analytical Chemistry 91, 8366–8373, doi:10.1021/acs.analchem.9b01178 (2019).

8 Cash, K. J. & Clark, H. A. Phosphorescent nanosensors for in vivo tracking of histamine levels. Anal Chem 85, 6312–6318, doi:10.1021/ac400575u (2013).

9 Barnea, G. et al. The genetic design of signaling cascades to record receptor activation. Proc Natl Acad Sci U S A 105, 64–69, doi:10.1073/pnas.0710487105 (2008).

10 Kroeze, W. K. et al. PRESTO-Tango as an open-source resource for interrogation of the druggable human GPCRome. Nat Struct Mol Biol 22, 362–369, doi:10.1038/nsmb.3014 (2015).

11 Jagadish, S., Barnea, G., Clandinin, T. R. & Axel, R. Identifying functional connections of the inner photoreceptors in Drosophila using Tango-Trace. Neuron 83, 630–644, doi:10.1016/j.neuron.2014.06.025 (2014).

12 Liu, Y. et al. Visualization of the activation of the histamine H3 receptor (H3R) using novel fluorescence resonance energy transfer biosensors and their potential application to the study of H3R pharmacology. FEBS J 285, 2319–2336, doi:10.1111/febs.14484 (2018).

13 Schihada, H. et al. Development of a Conformational Histamine H3 Receptor Biosensor for the Synchronous Screening of Agonists and Inverse Agonists. ACS Sens 5, 1734–1742, doi:10.1021/acssensors.0c00397 (2020).

14 Erdogmus, S. et al. Helix 8 is the essential structural motif of mechanosensitive GPCRs. Nat Commun 10, 5784, doi:10.1038/s41467-019-13722-0 (2019).

15 Jing, M. et al. A genetically encoded fluorescent acetylcholine indicator for in vitro and in vivo studies. Nat Biotechnol 36, 726–737, doi:10.1038/nbt.4184 (2018).

16 Sun, F. et al. A Genetically Encoded Fluorescent Sensor Enables Rapid and Specific Detection of Dopamine in Flies, Fish, and Mice. Cell 174, 481–496 e419, doi:10.1016/j.cell.2018.06.042 (2018).

17 Dong, A. et al. A fluorescent sensor for spatiotemporally resolved imaging of endocannabinoid dynamics in vivo. Nat Biotechnol 40, 787–798, doi:10.1038/s41587-021-01074-4 (2022).

18 Patriarchi, T. et al. Ultrafast neuronal imaging of dopamine dynamics with designed genetically encoded sensors. Science 360, doi:10.1126/science.aat4422 (2018).

19 Wu, Z., Lin, D. & Li, Y. Pushing the frontiers: tools for monitoring neurotransmitters and neuromodulators. Nat Rev Neurosci, doi:10.1038/s41583-022-00577-6 (2022).

20 Feng, J. et al. A Genetically Encoded Fluorescent Sensor for Rapid and Specific In Vivo Detection of Norepinephrine. Neuron 102, 745–761 e748, doi:10.1016/j.neuron.2019.02.037 (2019).

21 Conklin, B. R., Farfel, Z., Lustig, K. D., Julius, D. & Bourne, H. R. Substitution of three amino acids switches receptor specificity of Gq alpha to that of Gi alpha. Nature 363, 274–276, doi:10.1038/363274a0 (1993).

22 Inoue, A. et al. Illuminating G-Protein-Coupling Selectivity of GPCRs. Cell 177, 1933–1947 e1925, doi:10.1016/j.cell.2019.04.044 (2019).

23 Scammell, T. E., Jackson, A. C., Franks, N. P., Wisden, W. & Dauvilliers, Y. Histamine: neural circuits and new medications. Sleep 42, doi:10.1093/sleep/zsy183 (2019).

24 Inagaki, N. et al. Organization of histaminergic fibers in the rat brain. J Comp Neurol 273, 283–300, doi:10.1002/cne.902730302 (1988).

25 Saper, C. B., Scammell, T. E. & Lu, J. Hypothalamic regulation of sleep and circadian rhythms. Nature 437, 1257–1263, doi:10.1038/nature04284 (2005).

26 Strecker, R. E. et al. Extracellular histamine levels in the feline preoptic/anterior hypothalamic area during natural sleep-wakefulness and prolonged wakefulness: an in vivo microdialysis study. Neuroscience 113, 663–670, doi:10.1016/s0306-4522(02)00158-6 (2002).

27 Chu, M. et al. Extracellular histamine level in the frontal cortex is positively correlated with the amount of wakefulness in rats. Neurosci Res 49, 417–420, doi:10.1016/j.neures.2004.05.001 (2004).

28 Parmentier, R. et al. The brain H3-receptor as a novel therapeutic target for vigilance and sleep-wake disorders. Biochem Pharmacol 73, 1157–1171, doi:10.1016/j.bcp.2007.01.002 (2007).

29 Lohse, M. J. et al. Kinetics of G-protein-coupled receptor signals in intact cells. Brit J Pharmacol 153 Suppl 1, S125–132, doi:10.1038/sj.bjp.0707656 (2008).

30 Wan, J. et al. A genetically encoded sensor for measuring serotonin dynamics. Nat Neurosci 24, 746–752, doi:10.1038/s41593-021-00823-7 (2021).

31 Ligneau, X. et al. Neurochemical and behavioral effects of ciproxifan, a potent histamine H3-receptor antagonist. J Pharmacol Exp Ther 287, 658–666 (1998).

32 Gondard, E. et al. Enhanced histaminergic neurotransmission and sleep-wake alterations, a study in histamine H3-receptor knock-out mice. Neuropsychopharmacology 38, 1015–1031, doi:10.1038/npp.2012.266 (2013).

33 Ericson, H., Watanabe, T. & Kohler, C. Morphological analysis of the tuberomammillary nucleus in the rat brain: delineation of subgroups with antibody against L-histidine decarboxylase as a marker. J Comp Neurol 263, 1–24, doi:10.1002/cne.902630102 (1987).

34 Wada, H., Inagaki, N., Yamatodani, A. & Watanabe, T. Is the histaminergic neuron system a regulatory center for whole-brain activity? Trends in Neurosciences 14, 415–418 (1991).

35 Inagaki, N. et al. An analysis of histaminergic efferents of the tuberomammillary nucleus to the medial preoptic area and inferior colliculus of the rat. Exp Brain Res 80, 374–380, doi:10.1007/BF00228164 (1990).

36 Wenkai Lin et al. Whole-brain mapping of histaminergic circuit in mouse brain. Co-submitted (2022).

37 Zhou, F. M., Liang, Y. & Dani, J. A. Endogenous nicotinic cholinergic activity regulates dopamine release in the striatum. Nat Neurosci 4, 1224–1229, doi:10.1038/nn769 (2001).

38 Liu, C. et al. An action potential initiation mechanism in distal axons for the control of dopamine release. Science 375, 1378–1385, doi:10.1126/science.abn0532 (2022).

39 Panula, P. & Nuutinen, S. The histaminergic network in the brain: basic organization and role in disease. Nat Rev Neurosci 14, 472–487, doi:10.1038/nrn3526 (2013).

40 Ercan-Sencicek, A. G. et al. L-histidine decarboxylase and Tourette’s syndrome. N Engl J Med 362, 1901–1908, doi:10.1056/NEJMoa0907006 (2010).

41 Hardie, R. C. Is histamine a neurotransmitter in insect photoreceptors? J Comp Physiol A 161, 201–213, doi:10.1007/BF00615241 (1987).

42 Hardie, R. C. A histamine-activated chloride channel involved in neurotransmission at a photoreceptor synapse. Nature 339, 704–706, doi:10.1038/339704a0 (1989).

43 Barger, Z., Frye, C. G., Liu, D., Dan, Y. & Bouchard, K. E. Robust, automated sleep scoring by a compact neural network with distributional shift correction. PLoS One 14, e0224642, doi:10.1371/journal.pone.0224642 (2019).

44 Sun, F. et al. Next-generation GRAB sensors for monitoring dopaminergic activity in vivo. Nat Methods 17, 1156–1166, doi:10.1038/s41592-020-00981-9 (2020).

